# Remodelling of skeletal muscle myosin metabolic states in hibernating mammals

**DOI:** 10.1101/2023.11.14.566992

**Authors:** Christopher T. A. Lewis, Elise G. Melhedegaard, Marija M. Ognjanovic, Mathilde S. Olsen, Jenni Laitila, Robert A. E. Seaborne, Magnus Nørregaard Grønset, Chengxin Zhang, Hiroyuki Iwamoto, Anthony L. Hessel, Michel N. Kuehn, Carla Merino, Nuria Amigó, Ole Fröbert, Sylvain Giroud, James F. Staples, Anna V. Goropashnaya, Vadim B. Fedorov, Brian M. Barnes, Øivind Tøien, Kelly L. Drew, Ryan J. Sprenger, Julien Ochala

## Abstract

Hibernation is a period of metabolic suppression utilized by many small and large mammal species to survive during winter periods. As the underlying cellular and molecular mechanisms remain incompletely understood, our study aimed to determine whether skeletal muscle myosin and its metabolic efficiency undergo alterations during hibernation to optimize energy utilization. We isolated muscle fibers from small hibernators, *Ictidomys tridecemlineatus* and *Eliomys quercinus* and larger hibernators, *Ursus arctos* and *Ursus americanus*. We then conducted loaded Mant-ATP chase experiments alongside X-ray diffraction to measure resting myosin dynamics and its ATP demand. In parallel, we performed multiple proteomics analyses. Our results showed a preservation of myosin structure in *U. arctos* and *U. americanus* during hibernation, whilst in *I. tridecemlineatus* and *E. quercinus*, changes in myosin metabolic states during torpor unexpectedly led to higher levels in energy expenditure of type II, fast-twitch muscle fibers at ambient lab temperatures (20°C). Upon repeating loaded Mant-ATP chase experiments at 8°C (near the body temperature of torpid animals), we found that myosin ATP consumption in type II muscle fibers was reduced by 77-107% during torpor compared to active periods. Additionally, we observed Myh2 hyper-phosphorylation during torpor in *I. tridecemilineatus*, which was predicted to stabilize the myosin molecule. This may act as a potential molecular mechanism mitigating myosin-associated increases in skeletal muscle energy expenditure during periods of torpor in response to cold exposure. Altogether, we demonstrate that resting myosin is altered in hibernating mammals, contributing to significant changes to the ATP consumption of skeletal muscle. Additionally, we observe that it is further altered in response to cold exposure and highlight myosin as a potentially contributor to skeletal muscle non-shivering thermogenesis.

## Introduction

Hibernation is an adaptive strategy employed by many animals aiming to decrease their metabolic rate and improve survival, particularly during harsh, winter conditions where food supply is limited, and thermogenic demands are high [1]. During hibernation, mammals typically undergo a decrease in body temperature, heart, and breathing rates [2–4]. In so-called fat-storing hibernators, this is inherently accompanied by prolonged fasting and fatty acids become the main substrate for energy provision [5, 6]. Besides these common features associated with overall metabolic depression, there are also significant inter-species differences in the underlying strategies. For instance, small (< 8 kg) fat-storing hibernating mammals such as thirteen-lined ground squirrels (*Ictidomys tridecemlineatus*) or garden dormice (*Eliomys quercinus*) experience extended bouts of a hypo-metabolic state (torpor), punctuated by spontaneous periods of interbout euthermic arousals (IBA), during which metabolic activity transiently increases back to basal levels. During torpor, metabolic rate decreases below 5% of euthermic values and core body temperatures decrease from 35°C - 38°C to 4°C - 8°C [7, 8]. In contrast, either medium (10-20 kg, *e.g.* European badger, *Meles meles*) or large (> 20 kg, *e.g.* Eurasian brown bear, *Ursus arctos*, and American black bear, *Ursus americanus*) hibernating mammals exhibit a pronounced hypo-metabolic state (as low as 25% of their basal metabolic rate in the case of bears), but only experience a mild decline in body temperature (to 32-35°C depending on body size) that lasts for several winter months [1, 9, 10]. While species-specific physiological patterns are well-documented, the molecular and cellular mechanisms operative in individual organs to achieve these remain largely undefined.

Skeletal muscle constitutes approximately 45-55% of body mass and serves as a major determinant of basal metabolic rate and heat production [11, 12]. Previous studies have uncovered some specific metabolic changes in skeletal muscle during hibernation [6]. For example, a decrease in mitochondrial respiration, as well as a suppression of ATP production capacity, has previously been documented in skeletal muscles from *I. tridecemlineatus* during torpor [13]. Previous work has demonstrated that ground squirrels require an optimal dietary ratio of monounsaturated to polyunsaturated fats, approximately 2:1, during their fat-storing period to facilitate effective hibernation [14]. The proteome of *I. tridecemlineatus* has then been shown to be enriched for fatty acid β-oxidation during periods of torpor. However, some reliance upon carbohydrate metabolism is maintained. The activity of phosphoglucomutase (PGM1) is even increased during torpor [15]. In *U. arctos*, skeletal muscle exhibits a transition from carbohydrate utilization to lipid metabolism, coupled with a reduction in whole-tissue ATP turnover [16].

Muscle is organized into an array of fibers containing repeating sarcomeres, which are crucial for regulating not only contraction but also metabolism and thermogenesis [17]. Until recently, energy consumption in skeletal muscle was thought to be primarily linked to the activity of myosin during muscular contraction [17]. Additionally, thermogenesis in skeletal muscle was previously attributed primarily to the electron transport system, in some cases link to uncoupling of sarcoplasmic reticulum calcium ATPase (SERCA) [18, 19]. However, growing evidence that the sarcomeric metabolic rate and thermogenesis are also controlled by ‘relaxed’ myosin molecules is emerging [20]. Myosin heads in passive muscle (pCa > 8), can be in different resting metabolic states that maintain a basal level of ATP consumption. In the ‘disordered-relaxed’ (DRX) state, myosin heads are generally not bound to actin and structurally exist in a conformation (so-called ON state) where they primarily exist freely within the interfilamentous space in the sarcomere [21, 22]. In the ‘super-relaxed’ (SRX) state, myosin heads adopt a structural conformation against the thick filament backbone (so-called OFF state) [23]. This conformation sterically inhibits the ATPase site on the myosin head, significantly reducing both ATP turnover in these molecules and, therefore heat production. The SRX state has an ATP turnover rate five to ten times lower than that of myosin heads in the DRX state [24]. A 20% shift of myosin heads from SRX to DRX is predicted to increase whole-body energy expenditure by 16% and double skeletal muscle thermogenesis [24]. What remains to be determined is whether, across mammals, increasing the proportions of myosin in the DRX or SRX states serves as a physiological molecular mechanism to fine-tune metabolic demands and thermogenesis. This includes potential contributions to whole-body metabolic depression observed during hibernation.

Hence, in the present study, we hypothesized that a remodelling of the proportions of myosin DRX and SRX conformations within skeletal muscles occurs and is a major suppressor of ATP/metabolic demand during hibernation. A recent study on *I. tridecemlineatus* cardiac muscle supports this hypothesis, finding higher proportions of SRX during torpor [25]. We examined isolated skeletal myofibers extracted from both small and large hibernating mammals - *I. tridecemlineatus*, *E. quercinus*, *U. arctos,* and *U. americanus*. We employed a multifaceted approach: loaded Mant-ATP chase experiments to assess myosin conformation and ATP turnover time, X-ray diffraction for sarcomere structure evaluation, and proteomic analyses to quantify differential PTMs.

## Results

### Resting myosin metabolic states are preserved in skeletal muscles fibers of hibernating *Ursus arctos* and *Ursus americanus*

To investigate whether resting myosin DRX and SRX states and their respective ATP consumption rates were altered during hibernation, we utilized the loaded Mant-ATP chase assay in isolated permeabilized muscle fibers from *U. arctos* obtained during either summer (active period) or winter (hibernating period). A total of 104 myofibers were tested at ambient lab temperatures (20°C). A representative decay of the Mant-ATP fluorescence in single muscle fibers is shown, indicating ATP consumption by myosin heads (figure 1A). In both type I (myosin heavy chain - MyHC-I) and type II (MyHC-II) myofibers, we did not observe any change in the percentage of myosin heads in either the DRX (P1 in figure 1B) or SRX states (P2 in figure 1C). We also did not find any difference in their ATP turnover times as demonstrated by the preserved T1 (figure 1D) and T2 values (figure 1E). We calculated the ATP consumed by myosin molecules by using the following equation based on the assumption that the concentration of myosin heads within single muscle fibers is 220 μM [24]:

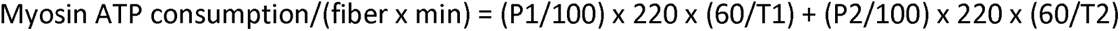

the ATP consumed by myosin molecules was not different between groups (figure 1F). We repeated these experiments on 95 myofibers obtained from summer and winter from *U. americanus*. A representative decay of the Mant-ATP fluorescence in single muscle fibers is shown (figure 1G). We found very similar results to the *U.* arctos that in both type I and type II muscle fibers, with no changes to resting myosin conformation, ATP turnover time or ATP consumption per fiber. These data indicate that myosin metabolic states are unchanged during hibernation in both *U. arctos* and *U. americanus*.

**Figure 1.**
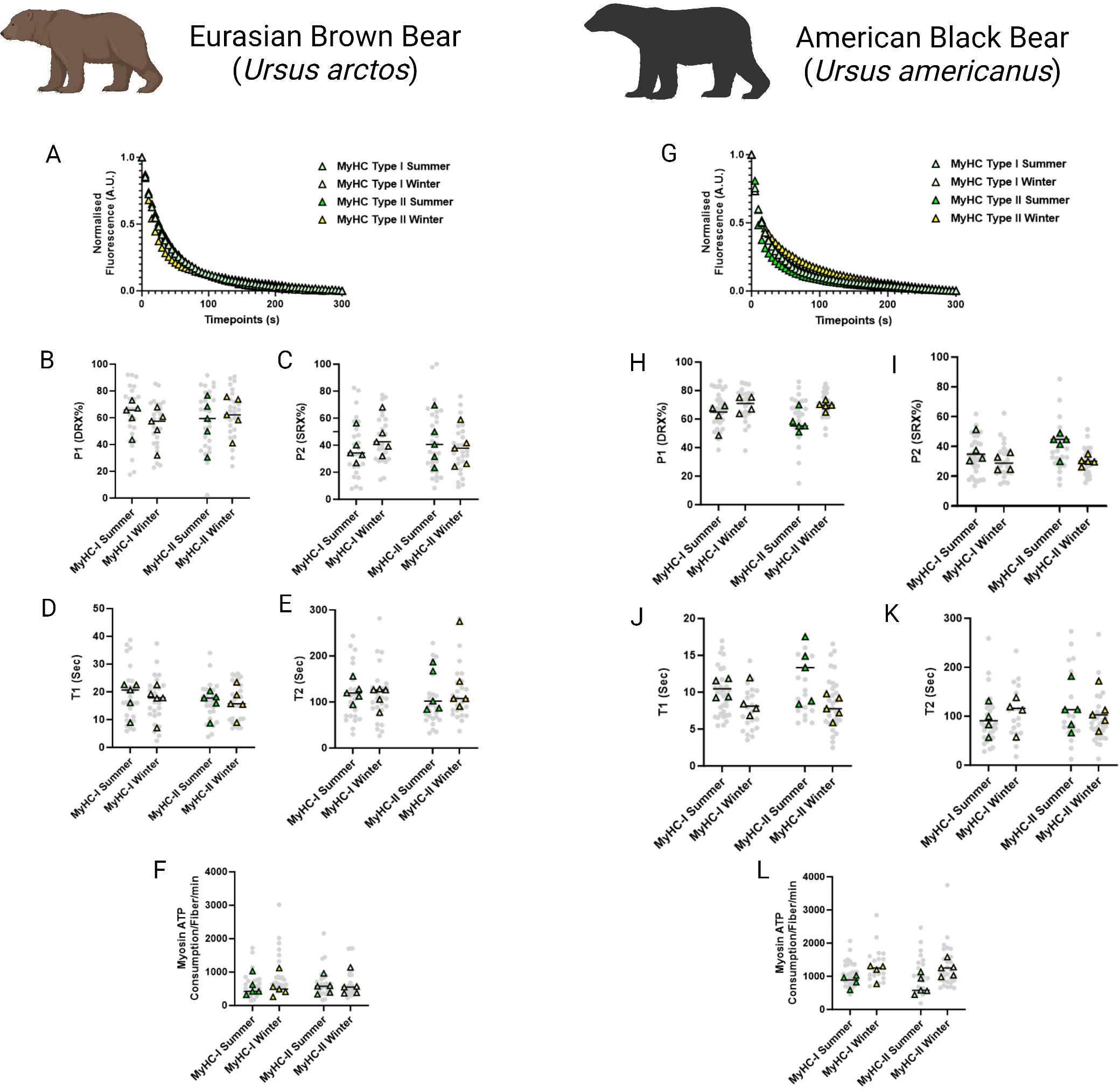
Myosin dynamics and myosin ATP consumption is unchanged in *Ursus arctos* and *Ursus americanus* during hibernation. **A.** Representative fluorescence mant-ATP decays from single muscle fibers isolated from *Ursus arctos* skeletal muscle measured over 300 seconds. **B-C.** Percentage of myosin heads in the P1/DRX (B) or P2/SRX (C) from *Ursus arctos* single muscle fibers obtained during summer (active) or winter (hibernating) periods. Values were separated based on each individual fiber was MyHC type I or MyHC type II. **D.** T1 value in seconds denoting the ATP turnover lifetime of the DRX. **E.** T2 value in seconds denoting the ATP turnover lifetime in seconds of the SRX. **F.** Calculated myosin ATP consumption values of each single muscle fiber per minute. This was calculated using the equation shown in the materials and methods section. **G.** Representative fluorescence mant-ATP decays from single muscle fibers isolated from *Ursus americanus* skeletal muscle measured over 300 seconds. **H-I.** Percentage of myosin heads in the P1/DRX (G) or P2/SRX (H) from *Ursus americanus* single muscle fibers obtained during summer (active) or winter (hibernating) periods. Values were separated based on each individual fiber was MyHC type I or MyHC type II. **J.** T1 value in seconds denoting the ATP turnover lifetime of the DRX. **K.** T2 value in seconds denoting the ATP turnover lifetime in seconds of the SRX. **L.** Calculated myosin ATP consumption values of each single muscle fiber per minute. Grey circles represent the values from each individual muscle fiber which was analyzed. Colored triangles represent the mean value from an individual animal, 8-12 fibers analyzed per animal. Statistical analysis was performed upon mean values. One-way ANOVA was used for statistical testing. n = 5 individual animals per group.

### Relaxed myosin metabolic states are disrupted in skeletal myofibers of small hibernators: *Ictidomys tridecemlineatus* and *Eliomys quercinus*

In smaller hibernators we utilized samples obtained from *E. quercinus* and *I. tridecemlineatus* during the summer active state (SA), IBA and torpor. In *E. quercinus*, a total of 146 myofibers were assessed at ambient lab temperatures. Consistent with *U. arctos* and *U. americanus*, we did not see any difference in the percentage of myosin heads in either the DRX (figure 2A) or SRX states (figure 2B). However, the ATP turnover time of myosin molecules in the DRX conformation (DRX T1) was 35% lower in IBA and torpor compared with SA in type I fibers and was 36% and 31% lower during IBA and torpor, respectively, compared to SA in type II fibers (figure 2C). The ATP turnover time of myosin heads in the SRX state (SRX T2) was 28% lower in in both IBA and torpor compared to SA for type I fibers and was lower in torpor by 26% compared with TA for type II fibers (figure 2D). All these changes were accompanied by a 56% greater myosin-based ATP consumption of in IBA compared to SA for type I fibers. In type II myofibers ATP consumption was 55% and 47% greater in IBA and torpor, respectively, compared with SA (figure 2E). In *I. tridecemlineatus*, a total of 156 muscle fibers were evaluated. In accordance with *E. quercinus,* we did not observe any modification in the percentage of myosin heads in the DRX conformation (P1) in the SRX conformation (P2) in either fiber type between SA, IBA or torpor (figure 2F, G). Nevertheless, DRX T1 was 35% lower for type I fibers in torpor compared to SA and was 29% lower in IBA and 46% in torpor compared to SA for type II fibers (figure 2H). SRX T2 was significantly lower in both IBA, and torpor compared to SA for type I by 49% and 29% respectively (figure 2I). SRX T2 was also significantly lower in type II fibers in both IBA, and torpor compared to SA by 31% and 27% respectively. Myosin-specific ATP consumption in type II muscle fibers during torpor was significantly higher at 99% compared to SA (figure 2J).

**Figure 2.**
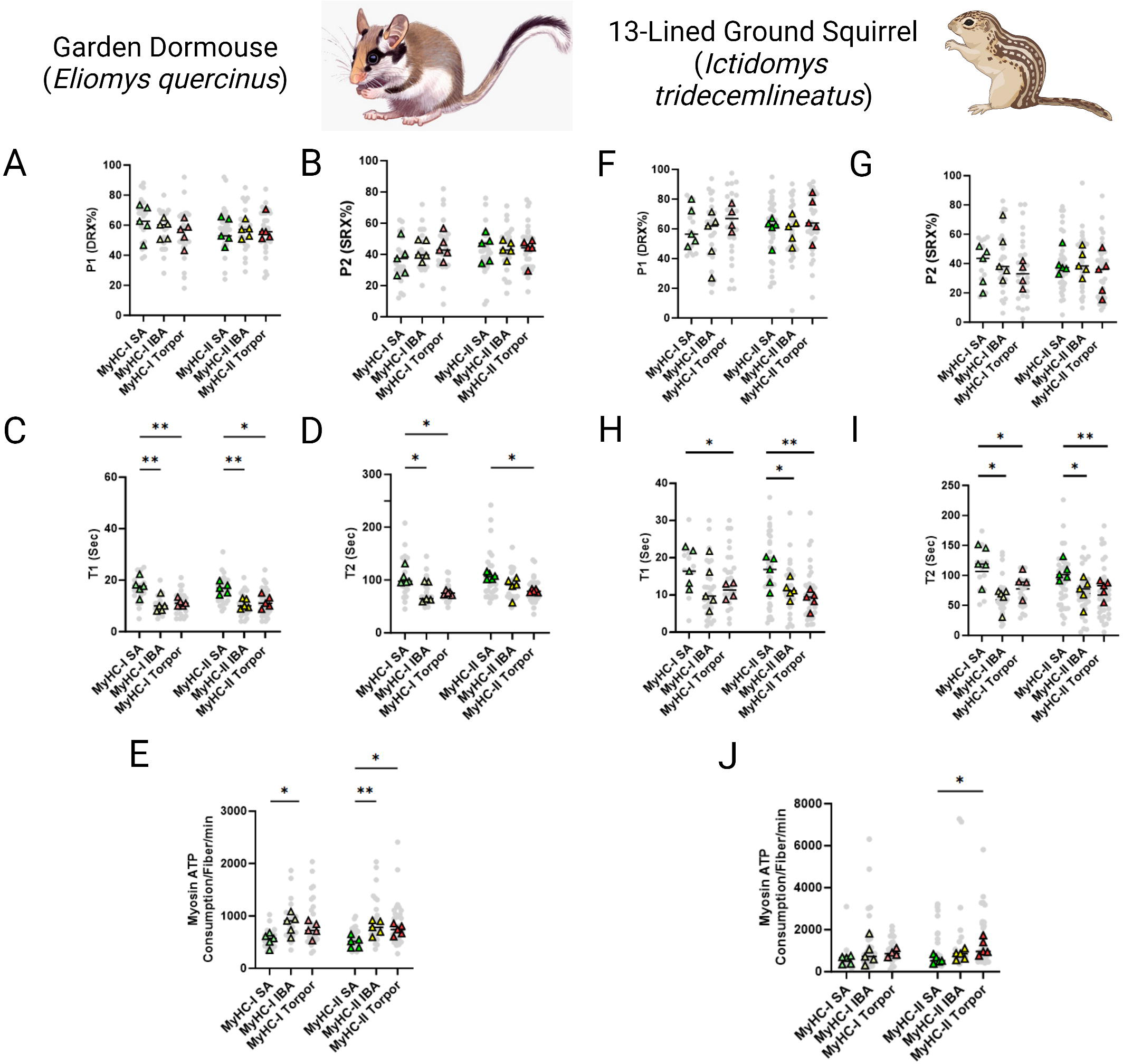
Myosin ATP turnover lifetime is reduced during hibernation in small hibernators, *Eliomys quercinus* & *Ictidomys tridecemlineatus*, resulting in an increase in myosin ATP consumption at ambient temperatures. **A-B.** Percentage of myosin heads in the P1/DRX (A) or P2/SRX (B) from *E. quercinus* single muscle fibers obtained during active, interbout arousal (IBA) or torpor periods. Values were separated based on each individual fiber was MyHC type I or MyHC type II. **C.** T1 value in seconds denoting the ATP turnover lifetime of the DRX in *E. quercinus*. **D.** T2 value in seconds denoting the ATP turnover lifetime in seconds of the SRX in *E. quercinus*. **E.** Calculated myosin ATP consumption values of each single muscle fiber per minute in *E. quercinus*. This was calculated using the equation shown in the materials and methods section. **F-G.** Percentage of myosin heads in the P1/DRX (F) or P2/SRX (G) from *Ictidomys tridecemlineatus* single muscle fibers obtained during summer active (SA), interbout arousal (IBA) or torpor periods. **H.** T1 value in seconds denoting the ATP turnover lifetime of the DRX in *Ictidomys tridecemlineatus*. **I.** T2 value in seconds denoting the ATP turnover lifetime in seconds of the SRX in *Ictidomys tridecemlineatus*. **J.** Calculated myosin ATP consumption values of each single muscle fiber per minute in *Ictidomys tridecemlineatus*. Grey circles represent the values from each individual muscle fiber which was analyzed. Colored triangles represent the mean value from an individual animal, 8-12 fibers analyzed per animal. Statistical analysis was performed upon mean values. One-way ANOVA was used to calculate statistical significance. * = *p* < 0.05, ** = *p* < 0.01. n = 5 individual animals per group.

To gain insights into the mechanisms that cause such unexpected myosin metabolic adaptive changes, we investigated whether these latter changes were accompanied by a structural alteration of myosin molecules. We collected and analysed X-ray diffraction patterns of thin muscle strips in *I. tridecemlineatus*. The ratio of intensities between the 1,0 and 1,1 reflections (I_1,1_ / I_1,0_; figure 3A) provides a quantification of myosin mass movement between thick and thin filaments. An increase in I_1,1_ / I_1,0_ reflects myosin head movement from thick to thin filaments, where increasing I_1,1_ / I_1,0 _ tracks myosin head movement from thick to thin filaments, signalling a transition from the OFF to ON state [26]. 1,0 and 1,1 equatorial intensities were quantified and the intensity ratio (I_1,1_ to I_1,0_) calculated. This intensity ratio was significantly lower in torpor compared to IBA (figure 3B), suggesting more myosin heads are structurally OFF in torpor vs IBA. A reorientation of the myosin heads between OFF and ON states can also be captured by the M3 reflection along the meridional axis of diffraction patterns (figure 3A). The M3 spacing represents the average distance between myosin crowns along the thick filament. An increase in M3 spacing signifies a reorientation of myosin heads from the OFF towards the ON states [27]. No differences were seen in M3 spacing or intensity (figure 3C, D), indicating that the orientation of myosin crowns along the thick filament are similar across all conditions (SA, IBA and torpor). Thick filament length, measured here by the spacing of the M6 reflection (figure 3A), was significantly greater in IBA compared to SA, and during torpor compared to both SA and IBA (figure 3E). Taken together we report a unique structural signature of muscle during hibernating periods in *I. tridecemlineatus*.

**Figure 3.**
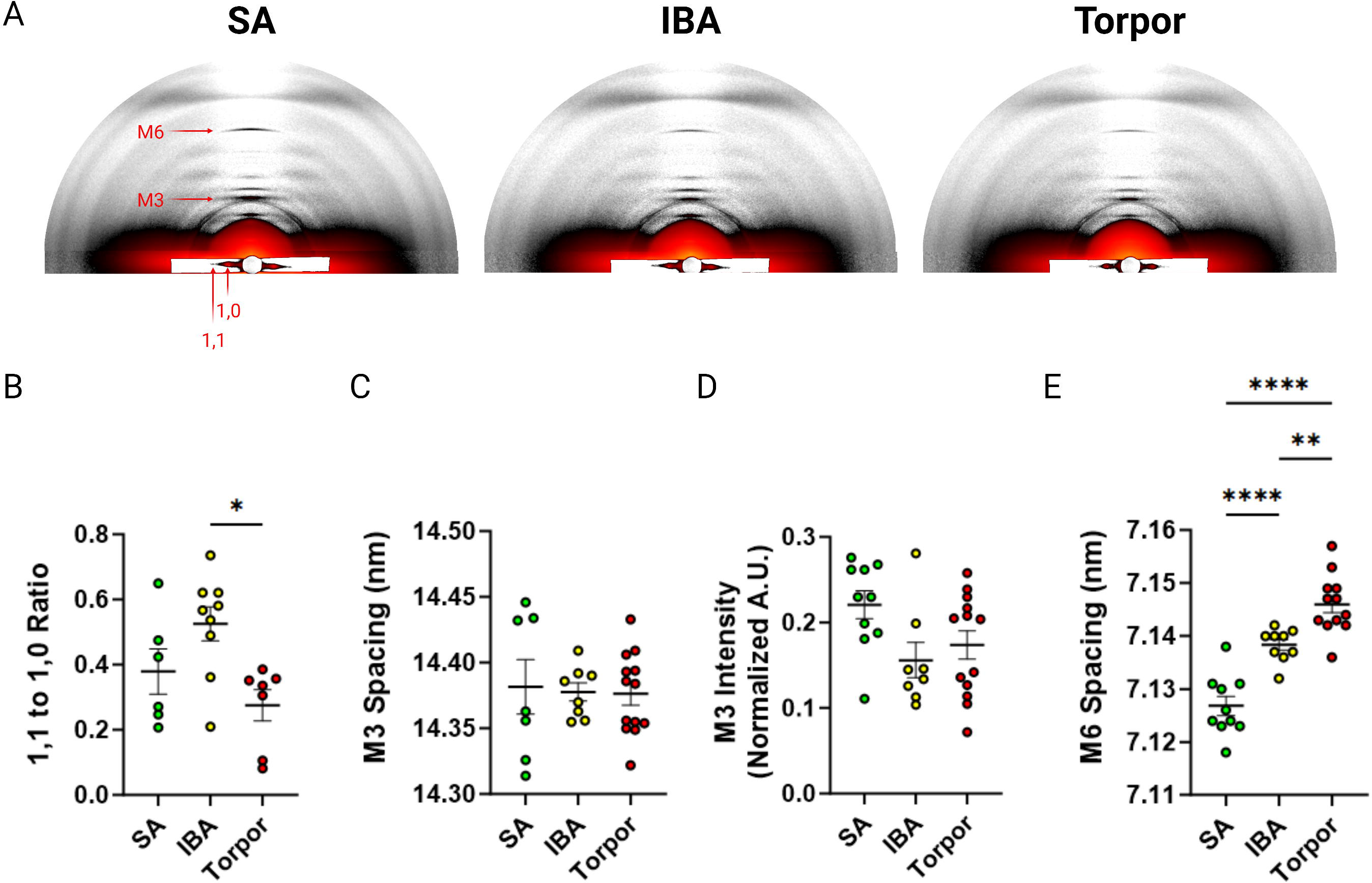
X-ray diffraction experiments of skeletal muscle from *Ictidomys tridecemlineaus* demonstrate changes in M6 myosin meridional spacing during torpor. **A.** Representative x-ray diffraction recordings from permeabilized skeletal muscle bundles from *Ictidomys tridecemlineatus* from summer active (SA), interbout arousal (IBA) and torpor. The M3 and M6 meridional reflections and the 1,0 and 1,1 equatorial reflections are indicated. **B.** Ratio of the 1,1 to 1,0 equatorial reflections from active, IBA and torpor skeletal muscle. **C.** M3 meridional spacing, measured in nm. **D.** Normalized intensity (A.U.) of the M3 meridional reflection. **E.** M6 meridional spacing, measured in nm. Colored circles represent the mean value obtained from each skeletal muscle bundle which was recorded. Data is displayed as mean ± SEM. One-way ANOVA was used to calculate statistical significance. * = *p* < 0.05, ** = *p* < 0.01, *** = *p* < 0.001. n = 5 individual animals per group.

### Myosin temperature sensitivity is lost in relaxed skeletal muscles fibers of hibernating *Ictidomys tridecemlineatus*

To mimic the drastic body temperature decrease experienced by small hibernators during torpor, we repeated the loaded Mant-ATP chase experiments in *I. tridecemlineatus* at 8°C. A total of 138 myofibers were assessed and similar to when measured at ambient lab temperatures, no changes to the conformation of myosin states were observed (supplementary figure 1). At these temperatures we observed that during torpor the DRX T1 was 77% compared to SA and 107% higher compared to IBA in type II muscle fibers (figure 4A). We further observed that during torpor the SRX T2 was 60% higher compared to SA and 64% higher compared to IBA (figure 4B). We then calculated the 20°C to 8°C degree ratios, allowing us to define myosin temperature sensitivity of DRX T1, SRX T2 and myosin ATP consumption (figure 4C, D and E). In SA and IBA, lowering the temperature led to a reduction in the myosin ATP turnover time of both the DRX and SRX, thus decreasing ATP consumption especially for type II muscle fibers. In contrast, during torpor lowering the temperature had opposite effects (figure 4C, D and E). This observation was particularly prominent in type II muscle fibers (figure 4C, D). From these results, we suggest that *I. tridecemlineatus* reduce their resting myosin ATP turnover rates in response to cold exposure to increase heat production via ATP hydrolysis.

**Figure 4.**
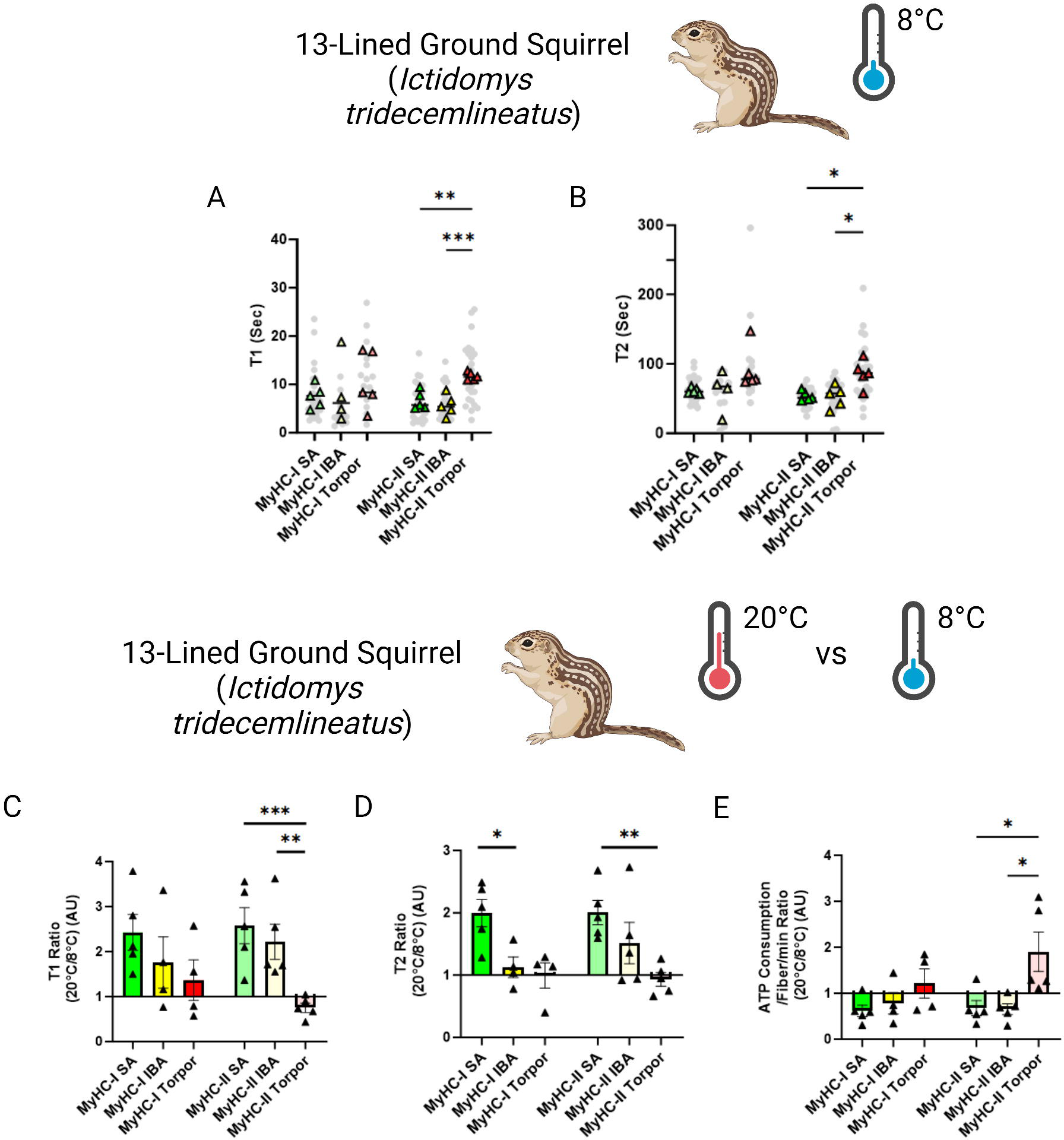
Myosin dynamics of *Ictidomys tridecemlineatus* are protected from temperature induced change during torpor, preventing an increase in myosin ATP consumption. **A.** T1 value in seconds denoting the ATP turnover lifetime of the DRX in *I. tridecemlineatus* at 8°C. **B.** T2 value in seconds denoting the ATP turnover lifetime in seconds of the SRX in *I. tridecemlineatus* at 8°C. **C.** Ratio of the T1 expressed as the mean value for each matched animal at 20°C/8°C, separated for fiber type. **D.** Ratio of the T2 expressed as the mean value for each matched animal at 20°C/8°C, separated for fiber type. **E.** Ratio of calculated myosin ATP consumption expressed as 20°C/8°C, separated for fiber type. Black triangles represent the mean ratio value for each animal. One-way ANOVA was used to calculate statistical significance. * = *p* < 0.05, ** = *p* < 0.01, *** = *p* < 0.001. n = 5 individual animals per group.

### Hyper-phosphorylation of Myh2 predictably stabilizes myosin backbone in hibernating *Ictidomys tridecemlineatus*

Based on our findings of discrepancies between the dynamics of myosin at different temperatures, we wanted to obtain a greater understanding of changes at the protein level during these different hibernating states. The myosin molecule is well-known to be heavily regulated by post-translational modifications (PTMs) [28–34]. We assessed whether hibernation impacts the level of phosphorylation and acetylation on the Myh7 and Myh2 proteins from *I. tridecemlineatus*. Myh2 exhibited significant differences in phosphorylation sites during torpor compared to SA and IBA (figure 5A). Three specific residues had significantly greater levels of phosphorylation: threonine 1039 (Thr1039-P), serine 1240 (Ser1240-P) and serine 1300 (Ser1300-P). These PTMs lie within the coiled-coil region of the myosin filament backbone (figure 5B, C). To define whether they have functional implications during hibernation, we utilized EvoEF, an *in-silico* programme which can characterize the effects of single amino acid residue substitutions on protein stability [35]. For our analysis, we replaced the three native residues where PTMs were found by aspartic acid (Asp), which chemically resembles phosphothreonine (Thr-P) and phosphoserine (Ser-P) [36]. Thr1039Asp, Ser1240Asp and Ser1300Asp had higher protein stabilities compared to Thr1039, Ser1240 and Ser1300, as attested by ΔΔG_Stability_ values greater than zero (figure 5D). The combination of these modifications does not counteract one another and thus are predicted to provide a high change in Myh2 stability (ΔΔG_Stability_ of 2.54, figure 5D).

**Figure 5.**
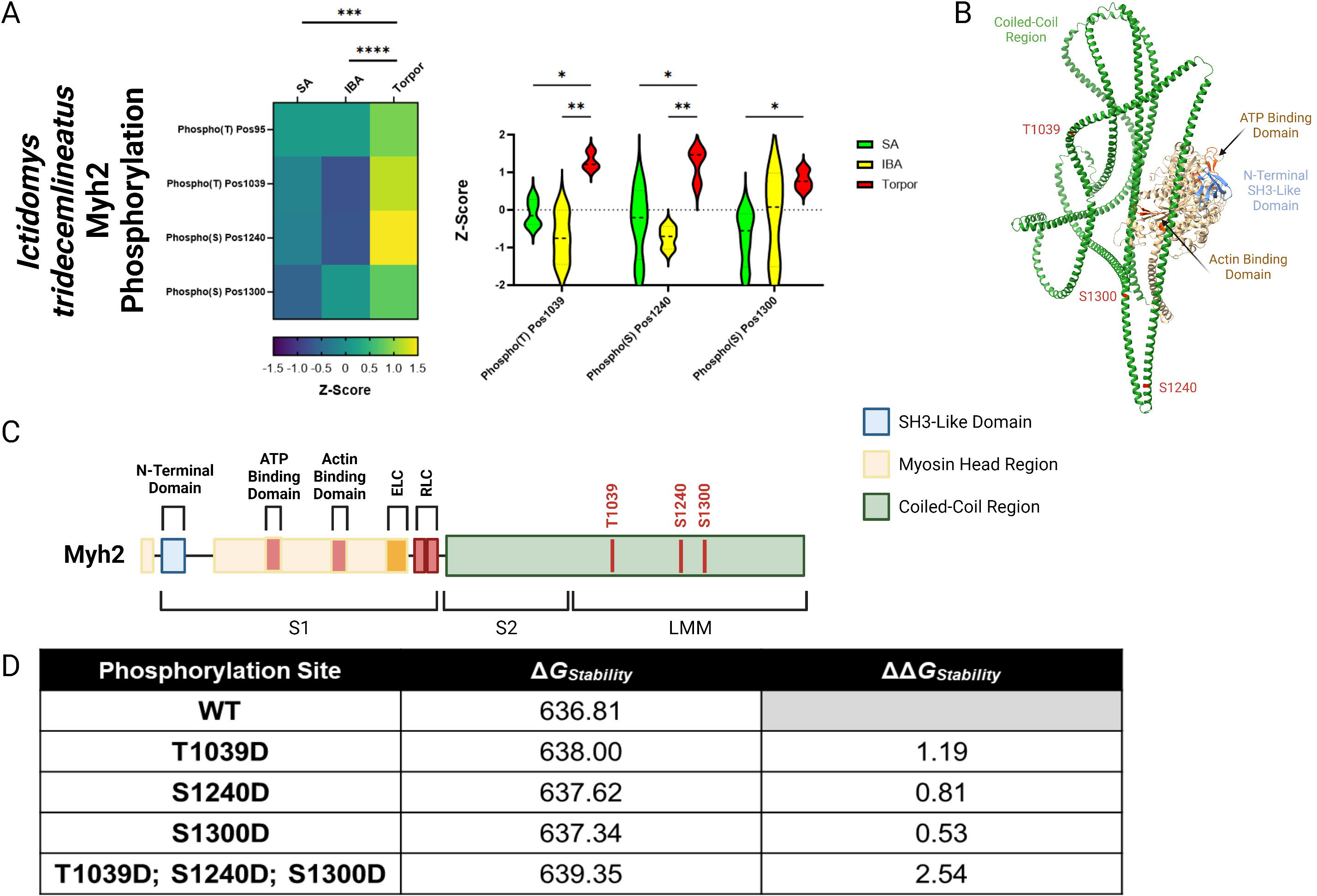
MYH2 protein in *Ictidomys tridecemlineatus* is hyper-phosphorylated during torpor, which is predicted to increase protein stability. **A.** Peptide mapping of differentiated phosphorylation sites upon MYH2 protein during SA, IBA and torpor periods. Heat map demonstrates all sites observed to be differentiated following the calculation of z-scores for each site. Z-scores > 0 equal hyper-phosphorylation and z-scores < 0 equal hypo-phosphorylation for each residue. Violin plot demonstrates significantly differentiated residues using z-scores. Two-way ANOVA with Šídák’s multiple comparisons test was used to calculate statistical significance. * = *p* < 0.05, ** = *p* < 0.01, *** = *p* < 0.001, **** = *p* < 0.0001. n = 5 individual animals per group. **B.** Chimera of MYH2 protein created using ChimeraX software. Important regions of the protein are annotated including coiled-coil region, ATP binding domain, actin binding domain and N-terminal SH3-like domain. Also, significantly hyper-phosphorylated residues are highlighted in red. **C.** Schematic of MYH2 protein with regions and hyper-phosphorylated resides annotated in red. Figure made in BioRender. **D.** EvoEF calculations of protein stability in both wild type and phosphor-mimetic mutants. Aspartic acid was used to mimic phospho-threonine/phospho-serine due to their chemical similarity. ΔG_Stability_ indicates the stability score for the protein in its corresponding configuration. ΔΔG_Stability_ represents the change in stability in mutant proteins versus the wild type protein. ΔΔG_Stability_ of > 0 represents an increase in the stability of a mutant versus wild type.

We did not identify any hibernation-related changes in Myh7 phosphorylation or acetylation, nor did we detect any alterations in Myh2 acetylation in *I. tridecemlineatus*.

To validate the significance of the PTM findings, a parallel PTM analysis was performed for *U. arctos*, a species in which myosin metabolic states are unaffected by hibernation (figure 1). In this context, minimal changes in phosphorylated or acetylated residues were observed in the Myh7 and Myh2 proteins (supplementary figures 2 and 3). These minimal modifications occurred on amino acids distinct from those in *I. tridecemlineatus* (supplementary figures 2 and 3). Overall, our PTM analyses and related simulations indicate that torpor is associated with a Myh2 hyper-phosphorylation possibly impacting myosin filament backbone stability in *I. tridecemlineatus*.

### Sarcomeric proteins are dysregulated in resting skeletal myofibers of hibernating *Ictidomys tridecemlineatus*

In addition to PTMs, myosin binding partners and/or surrounding proteins may change during torpor and may contribute to disruptions of myosin metabolic states in *I. tridecemlineatus*. We performed an untargeted global proteomics analysis on isolated muscle fibers. Principal component analyses showed that whilst SA muscle fibers form a separate entity, both IBA and torpor myofibers are clustered, suggesting a common proteomics signature during the two states of hibernation (figure 6A). Differentially expressed proteins included molecules involved in sarcomere organisation and function (e.g., SRX-determining MYBPC2 and sarcomeric scaffold-defining ACTN3) as well as molecules belonging to metabolic pathways (e.g., lipid metabolism-related HMGCS2, carbohydrate metabolism-linked PDK4) (Figure 6B, 6C as well as supplementary figures 4 and 5). Gene ontology analyses including the top five up-/downregulated proteome clusters reinforced the initial findings. They emphasized proteins related to ‘muscle system process’, ‘fiber organization’, ‘muscular contraction’ or ‘muscle development’ and to ‘lipid or carbohydrate metabolism’ pathways (figure 6D-G). To complement these results, we performed profiling of core metabolite and lipid contents within the myofibers of *I. tridecemlineatus*. Lipid measurements were significantly lower during IBA, and torpor compared to SA (e.g., omega-6 and omega-7 fatty acids - supplementary figure 6), in line with previous studies [37].

**Figure 6.**
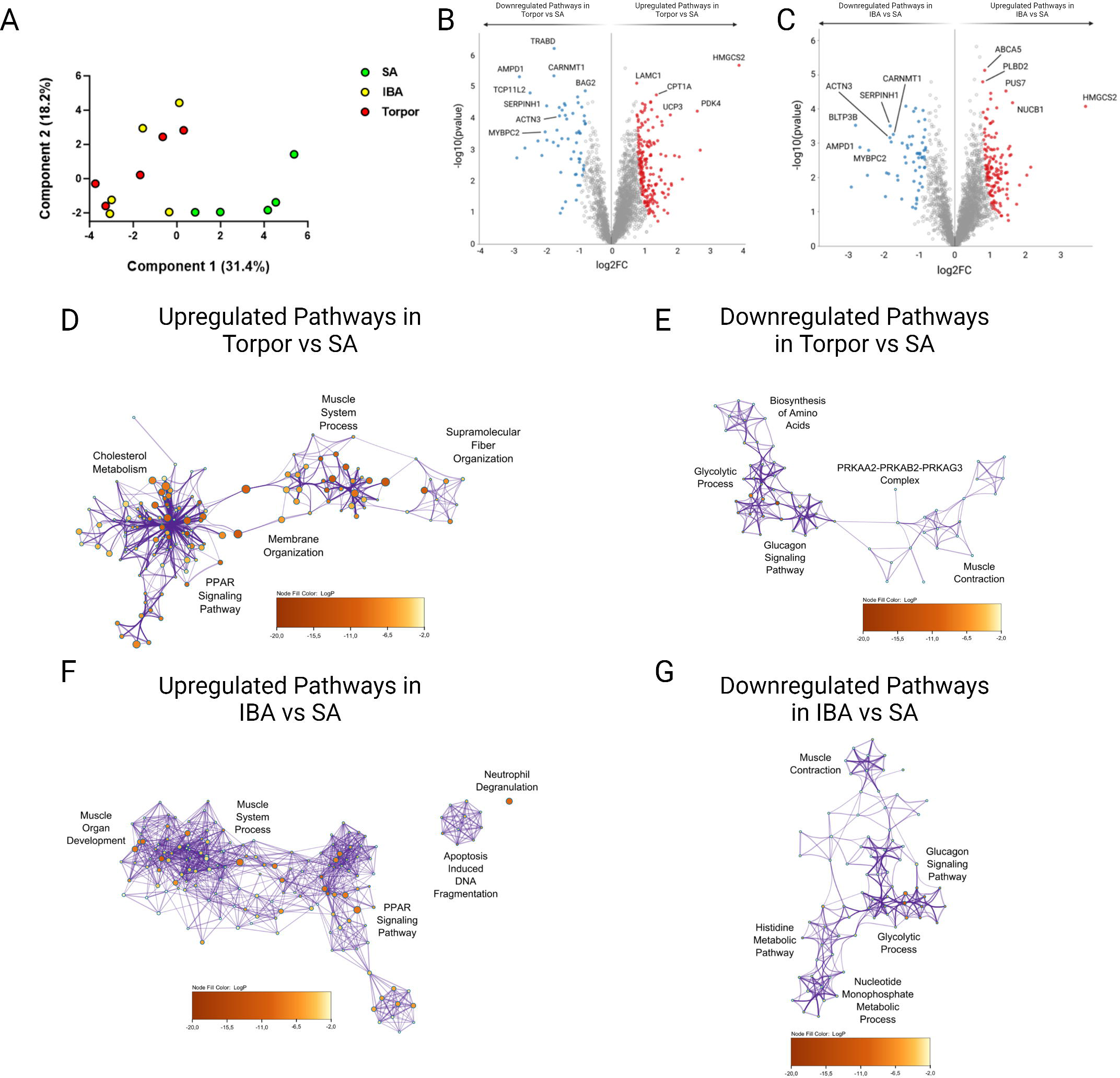
Global proteome analysis demonstrates changes to metabolic and sarcomeric changes in skeletal muscle fibers from *Ictidomys tridecemlineatus* during IBA and torpor. **A.** Principal component analysis for all animals analyzed during SA, IBA and torpor periods. **B.** Volcano plot displaying proteins which are differentially expressed during torpor vs active periods. FDR < 0.01. Red circles are upregulated proteins and blue circles are downregulated proteins. Highly differentiated proteins of interest are annotated with their respective protein name. **C.** Volcano plot displaying proteins which are differentially expressed during IBA vs SA periods. Red circles are upregulated proteins and blue circles are downregulated proteins. Highly differentiated proteins of interest are annotated with their respective protein name. **D.** Ontological associations between proteins upregulated during torpor vs SA periods. The top five association clusters are annotated on the network. A full list of clusters and the proteins lists included in clusters are available in supplementary figure 1 and supplementary table 2. **E.** Ontological associations between proteins downregulated during torpor vs SA periods. The top five association clusters are annotated on the network. A full list of clusters and the proteins lists included in clusters are available in supplementary figure 1 and supplementary table 2. **F.** Ontological associations between proteins upregulated during IBA vs SA periods. The top five association clusters are annotated on the network. A full list of clusters and the proteins lists included in clusters are available in supplementary figure 2 and supplementary table 3. **G.** Ontological associations between proteins downregulated during IBA vs SA periods. The top five association clusters are annotated on the network. A full list of clusters and the proteins lists included in clusters are available in supplementary figure 2 and supplementary table 3. Gene ontology networks were established using Metascape and visualized using Cytoscape. Detailed information upon the statistical testing used is available in the methods section. FDR < 0.01 significantly differentially expressed proteins were used to establish networks. Purple lines indicate a direct interaction. Circle size is determined by enrichment and color is determined by *p* value. n = 5 individual animals per group.

Untargeted global proteomics analysis was performed in *U. arctos* where myosin metabolic states are preserved during hibernation (Figure 1). As for *I. tridecemlineatus*, the principal component analysis emphasized differences between active and hibernating *U. arctos* (supplementary figure 7). Differentially expressed proteins, complemented by gene ontology analyses, underscored hibernation-related shifts in metabolic proteins as in *I. tridecemlineatus* (supplementary figures 7 and 8). However, sarcomeric proteins did not appear as differentially expressed molecules, highlighting their potential contribution to the adaptation of myosin in hibernating *I. tridecemlineatus*.

## Discussion

In the present study, our objective was to investigate whether modulating muscle myosin DRX and SRX states could serve as a key mechanism for reducing ATP/metabolic demand during mammalian hibernation. Contrary to our hypothesis, our results indicate that during hibernation small mammals such as *I. tridecemlineatus* or *E. quercinus* modulate their myosin metabolic states unexpectedly by increasing energy expenditure of sarcomeres at ambient temperatures. At 8°C, muscle fibers from *I. tridecemlineatus* obtained during SA and IBA phases displayed a significant rise in myosin-based ATP consumption. Conversely, fibers sampled during torpor bouts did not exhibit this cold-induced increase. These data suggest that small hibernators may stabilize myosin during torpor to prevent cold-induced increases in energy expenditure and thus increased heat production. Overall, our results also demonstrate a preferential adaptation of type II, fast-twitch, muscle fibers. Type II muscle fibers generally have more plasticity than type I, slow-twitch, fibers and have been demonstrated to undergo behavioural and fiber type transition in response to both metabolic and exercise stimuli [38–41].

### Resting Myosin Conformation is Unchanged During Hibernation

In contrast to our study hypothesis, we demonstrated that the animals studied continue to maintain their active levels of myosin in the more metabolically active disordered-relaxed state (DRX) during periods of metabolic shutdown. This is of particular interest when compared to a previous study by Toepfer *et al.,* who observed that in cardiac muscle from *I. tridecemlineatus* the percentage of myosin heads in the DRX conformation was lower in periods of torpor vs SA and IBA [25]. Maintenance of the resting myosin conformation to active levels during hibernating periods may be to prevent the onset significant muscular atrophy during hibernation. Hibernating animals have evolved mechanisms that prevent skeletal muscle atrophy during the extended periods of immobilization inherent to hibernation [15, 42–44]. Further research into changes to myosin head conformation in human atrophy and immobilization models would provide an interesting comparison to these data and potentially highlight resting myosin conformation as a novel target in the treatment of sarcopenia associated with aging and/or inactivity. It would furthermore ideally be possible to increase the biological sample size of all the species analysed in this study to further confirm the results which we report, particularly as modest differences are seen in the resting myosin conformation values of *U. americanus*.

### Resting myosin ATP consumption is higher during hibernation in small mammals at ambient temperature

Our findings demonstrate that in small hibernators such as *I. tridecemlineatus* and *E. quercinus*, the ATP turnover time of relaxed myosin molecules (in both DRX and SRX conformations) is faster during torpor (and IBA), especially in type II muscle fibers, leading to an unexpected overall increased ATP consumption. Accordingly, a few studies investigating human pathological conditions have reported disruptions of the myosin ATP turnover times in resting isolated skeletal myofibers, but their actual impacts have never been thoroughly investigated [45–47]. Here, originally, we estimated the consequences on the actual energy consumption of sarcomeres/muscle fibers. Our results of higher ATP consumption during torpor (and IBA) could, at first glance, be seen as counter-intuitive but is an indication of adaptation to the myosin protein during hibernating periods. It was therefore essential to investigate if these changes were also observed at a lower temperature, more relevant to the actual temperature of skeletal muscle in small hibernators during hibernation [48].

### Resting myosin ATP turnover time is protected from cold-induced change during torpor in *Ictidomys Tridecemlineatus*

A critical difference between the large hibernators, *U.* arctos and *U. americanus*, and the small hibernators, *E. quercinus* and *I. tridecemlineatus*, during their hibernation periods is core body temperature. Whilst the large hibernators only undergo a modest temperature decrease during hibernation, small hibernators reduce their core body temperature drastically to between 4°C - 8°C [48–50]. We repeated the Mant-ATP chase assays at 8°C to mimic the environment of physiological torpor. Interestingly, lowering the temperature decreased DRX and SRX-linked ATP turnover times during active periods (in SA and IBA), especially in type II myofibers from *I. tridecemlineatus*, inducing an increase in ATP consumption. Metabolic organs such as skeletal muscle are well-known to increase core body temperature in response to significant cold exposure (i) by inducing rapid involuntary contractions known as shivering [51, 52] or (ii) a process named non-shivering thermogenesis (NST). NST has traditionally been attributed to processes in brown adipose tissue, however, in recent years, skeletal muscle has also been shown to contribute to heat production via NST [53]. NST in skeletal muscle is stimulated by Ca^2+^-slippage by the sarcoplasmic reticulum Ca^2+^-ATPase (SERCA) in a cascade of molecular events controlled by a protein called sarcolipin [54–57]. Mammals have evolved a mechanism of resistance to ryanodine receptor (RyR) opening via rises in Ca^2+^ to allow for Ca^2+^ leak which is activated by cAMP via the β-adrenergic system [58, 59]. This Ca^2+^ slippage leads to the uncoupling of SERCA activity from Ca^2+^ transport across the sarcoplasmic reticulum. Consequently, ATP hydrolysis does not fuel ion transport. Instead, the resultant ADP stimulates heat production via the mitochondrial electron transport system [57, 60]. Here we propose that, in addition to SERCA, myosin also contributes to NST in small hibernators. Interestingly, our group has previously demonstrated that the resting myosin dynamics are altered in patients with RYR1 mutation-related myopathies[47].

Of potential further interest to the regulation of myosin would be the differential expression of heat shock proteins (HSPs) during hibernation. Various HSPs have been observed to be differentially expressed during hibernation in mammals such as bears and bats[61, 62]. This is of relevance to the data from our study as HSPs have been demonstrated to be able to bind sarcomeric proteins and regulate their turnover, including that of myosin itself [63, 64]. The proteins they have been shown to interact with include the cardiac isoform of myosin binding protein-C an important regulator of resting myosin conformation in the heart [65].

As the biopsies which were used in this study were all obtained from the hind leg of the animals studied, it is important to consider that the myosin dynamics may differ if these biopsies were sampled from different areas in the body. This is particularly important different areas of the body can have different core temperature and in some distal muscles, shivering does not occur[66]. A study from Aydin *et al.,* demonstrated in mice that when a shivering muscle, soleus, was prevented from undergoing non-shivering thermogenesis via knock-out of UCP1 and were subsequently exposed to cold temperatures, the force production of these muscles was significantly reduced due to prolonged shivering [66]. These results do suggest that even in shivering muscle, non-shivering thermogenesis plays a key role in the generation of heat for survival and for the maintenance of muscle performance. Further work examining potential differences in the resting myosin dynamics of muscles sampled from different sites of the body would be of importance to the field in the future.

Essential to our findings were the simultaneous observations that these cold induced changes in myosin ATP turnover times in each resting myosin state were not observed in *I. tridecemlineatus* samples obtained during torpor. *I. tridecemlineatus* and other similar small hibernators require a significant reduction in their body temperature to survive during winter periods [48, 67]. Therefore, the inhibition of excess heat production via myosin ATP hydrolysis is likely a protective mechanism which has evolved to facilitate reductions in core body temperature and wider metabolic shutdown during torpor.

### Myh2 is hyper-phosphorylated during torpor increasing protein stability in *Ictidomys tridecemlineatus*

The exact causes of all the above alterations remain unclear but may be linked to unusual PTMs directly targeting MyHCs. Thus, hyper-phosphorylation stabilizing the myosin filament backbone of the Myh2 protein through Thr1039-P, Ser1240-P and Ser1300-P during torpor is proposed as a main potential underlying biophysical mechanism. Interestingly, *in silico* molecular dynamics simulations mimicking close-by phosphorylations (Thr1309-P and Ser1362-P) have previously demonstrated a structural impairment of the myosin filament backbone [47]. This is consistent with our X-ray diffraction experiments where M6 spacing was found to be greater during torpor and is indicative of a unique structural configuration of the thick filament during hibernation [26]. Besides PTMs, another potential cause of the myosin metabolic remodelling may be changes to myofibrillar protein expression. Our global untargeted proteomics analysis reveals that *I. tridecemlineatus* undergo a subtle reorganization in sarcomeric protein content [15, 16, 42]. Our results are consistent with previous similar analysis on *I. tridecemlineatus* from Hindle *et. al.,* who identified significant changes in carbohydrate metabolism but also changes in sarcomere and cytoskeletal organisation in SA vs torpor muscle [15]. In our analysis we observed examples of type II muscle fiber specific proteins which were highly differentially expressed. ACTN3 (α-actinin-3), a type II fiber-specific molecule involved in linking adjacent sarcomeres, was notably downregulated during hibernation [68]. Its depression in mammals unexpectedly appears to confer superior cold resistance and heat generation in skeletal muscle [68–70]. Another type II fiber-specific protein significantly downregulated in both torpor and IBA was MYBPC2 (fast skeletal myosin binding protein-C) [65, 71]. Its loss in skeletal mammals of rodents is thought to directly interfere with myosin conformation [71]. Taken together, we believe that the aberrant PTMs and/or protein expression remodelling do play a role in modifying the myosin filament stability and this all may be triggered by the well-known decreased tension on sarcomeres during hibernation [72, 73].

## Conclusion

Our findings suggest ATP turnover adaptations in DRX and SRX myosin states occur in small hibernators like *I. tridecemlineatus* during hibernation and cold exposure. In contrast, larger mammals like *U. arctos* and *U. americanus* show no such changes, likely due to their stable body temperature during hibernation. This supports our hypothesis that myosin serves as a non-shivering thermogenesis regulator in mammals, a mechanism inhibited during torpor.

## Methods

### Samples collection and cryo-preservation

Gastrocnemius muscles from *I. tridecemlineatus* and *E. quercinus* were collected from animals during the summer (when subjects cannot hibernate) as well as from those approximately half-way through a torpor bout and during IBA. Animal husbandry and hibernation status monitoring (using temperature telemetry or biologging) are described elsewhere [74–76]. Tissues were excised and frozen immediately in liquid N_2_, and subsequently stored at -80°C. Muscles were shipped from Canada (*I. tridecemlineatus*) or Austria (*E. quercinus*) to Denmark on dry ice. For *E. quercinus*, all procedures have been discussed and approved by the institutional ethics and animal welfare committee in accordance with GSP guidelines and national legislation (ETK-046/03/2020, ETK-108/06/2022), and the national authority according to §§29 of Animal Experiments Act, Tierversuchsgesetz 2012 - TVG 2012 (BMBWF-68.205/0175-V/3b/2018). For *I. tridecemlineatus* all procedures were approved by the Animal Care Committee at the University of Western Ontario and conformed to the guidelines from the Canadian Council on Animal Care.

In Dalarna, Sweden, subadult (2.5 years to 5.5 years) Scandinavian brown bears (*U.* arctos) were sedated during hibernation and active periods as part of the Scandinavian Brown Bear Project. Each bear was outfitted with GPS collars and VHF transmitters, enabling location tracking in dens during winter and in natural habitats during active months. Bears were located in their dens in late February and again, from a helicopter, in late June. For winter sedation, a cocktail of medetomidine, zolazepam, tiletamine, and ketamine was used. During summer captures, bears were sedated from a helicopter using a higher dose of medetomidine, zolazepam, and tiletamine, omitting ketamine, to adjust for increased metabolic activity [77]. All experiments on brown bears were performed with approval by the Swedish Ethical Committee on Animal Research (C18/15 and C3/16).

Protocols for black bear experiments were approved by the University of Alaska Fairbanks, Institutional Animal Care and Use Committee (IACUC nos. 02-39, 02-44, 05-55, and 05-56). Animal work was carried out in compliance with the IACUC protocols and ARRIVE guidelines.

Animal care, monitoring of physiological conditions of the black bear (*U. americanus*) and tissue harvesting were described previously [78]. Before tissue sampling, bears (51 – 143 kg) were captured in the field by Alaska Department of Fish and Game in May–July and kept in an outdoor enclosure for at least 2 months to allow adaptation for changes in mobility. Feeding was stopped 24 h before summer active animals were euthanized. Hibernating bears were without food or water since October 27 and euthanized for tissue sampling between March 1 and 26, about 1 month before expected emergence from hibernation. Core body temperature was recorded with radio telemetry and oxygen consumption and respiratory quotient were monitored in hibernating bears with open flow respirometry. Samples of quadriceps muscle have been collected from captive hibernating and summer active males older than 2 years and banked at -80°C.

### Solutions

As previously published [79, 80], the relaxing solution contained 4 mM Mg-ATP, 1 mM free Mg^2+^, 10^-6^ mM free Ca^2+^, 20 mM imidazole, 7 mM EGTA, 14.5 mM creatine phosphate and KCl to adjust the ionic strength to 180 mM and pH to 7.0. Additionally, the rigor buffer for Mant-ATP chase experiments contained 120 mM K acetate, 5 mM Mg acetate, 2.5 mM K_2_HPO_4_, 50 mM MOPS, 2 mM DTT with a pH of 6.8.

### Muscle preparation and fibre permeabilization

Cryopreserved muscle samples were dissected into small sections and immersed in a membrane-permeabilising solution (relaxing solution containing glycerol; 50:50 v/v) for 24 hours at -20°C, after which they were transferred to 4°C. These bundles were kept in the membrane-permeabilising solution at 4°C for an additional 24 hours. After these steps, bundles were stored in the same buffer at -20°C for use up to one week [81, 82].

### Mant-ATP chase experiments

On the day of the experiments, bundles were transferred to the relaxing solution and individual muscle fibres were isolated. Their ends were individually clamped to half-split copper meshes designed for electron microscopy (SPI G100 2010C-XA, width, 3 mm), which had been glued to glass slides (Academy, 26 x 76 mm, thickness 1.00-1.20 mm). Cover slips were then attached to the top to create a flow chamber (Menzel-Gläser, 22 x 22 mm, thickness 0.13-0.16 mm) [79, 80]. Subsequently, at 20°C, myofibers with a sarcomere length of 2.00 µm were kept (assessed using the brightfield mode of a Zeiss Axio Scope A1 microscope). Each muscle fibre was first incubated for five minutes with a rigor buffer. A solution containing the rigor buffer with added 250 μM Mant-ATP was then flushed and kept in the chamber for five minutes. At the end of this step, another solution made of the rigor buffer with 4 mM ATP was added with simultaneous acquisition of the Mant-ATP chase.

For fluorescence acquisition, a Zeiss Axio Scope A1 microscope was used with a Plan-Apochromat 20x/0.8 objective and a Zeiss AxioCam ICm 1 camera. Frames were acquired every five seconds with a 20 ms acquisition/exposure time using at 385nm, for five minutes. Three regions of each individual myofiber were sampled for fluorescence decay using the ROI manager in ImageJ as previously published [79, 80]. The mean background fluorescence intensity was subtracted from the average of the fibre fluorescence intensity for each image. Each time point was then normalized by the fluorescence intensity of the final Mant-ATP image before washout (T = 0). These data were then fit to an unconstrained double exponential decay using Graphpad Prism 9.0:

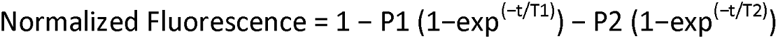

where P1 (DRX) is the amplitude of the initial rapid decay approximating the disordered-relaxed state with T1 as the time constant for this decay. P2 (SRX) is the slower second decay approximating the proportion of myosin heads in the super-relaxed state with its associated time constant T2 [79]. Mant-ATP Chase Experiment were performed at ambient lab temperature (20°C) for all samples unless otherwise stated. An additional setup was made to do the experiments at colder temperatures (∼8 °C). All slides were prepared in the same manner as mentioned above until incubation with rigor buffer and Mant-ATP buffer. All slides were kept on a metal plate on ice while incubated with both rigor and Mant-ATP buffer. The temperature of the chambers was measured (∼8 °C) before flushing with ice-cold ATP.

### Fiber-type staining

Fiber typing After the completion of the Mant-ATP chase experiments, individual fibers were stained with an anti-MyHC slow/type I antibody (A4.951; IgM isoform: 1:50, DSHB). Fibers were then washed in PBS and incubated with a secondary antibody conjugated to Alexa 647 in a goat serum (Thermo Fisher Scientific, dilution 1:1,000). After washing, the muscle fibers were mounted in Fluoromount, and images were taken with a Zeiss AXIO Lab A1 microscope (Carl Zeiss AG, GE, objectives ×20 and ×10). Positive staining with the MyHC β-slow/type I antibody indicated a type I muscle fiber and negative staining with the MyHC β-slow/type I antibody indicated a type II muscle fiber. Comparisons between muscle fibers sampled in summer or winter were then separated accordingly. Fiber-type breakdown and analysis for all samples used in this study are shown in Supplementary Table 1.

### X-ray diffraction recordings and analyses

Thin muscle bundles were mounted and transferred to a specimen chamber which was filled with the relaxing buffer. The ends of these thin muscle bundles were then clamped at a sarcomere length of 2.00 µm. Subsequently, X-ray diffraction patterns were recorded at 15°C using a CMOS camera (Model C11440-22CU, Hamamatsu Photonics, Japan, 2048 x 2048 pixels) in combination with a 4-inch image intensifier (Model V7739PMOD, Hamamatsu Photonics, Japan). The X-ray wavelength was 0.10 nm and the specimen-to-detector distance was 2.14 m. For each preparation, approximately 20-50 diffraction patterns were recorded at the BL40XU beamline of SPring-8 and were analysed as described previously [83]. To minimize radiation damage, the exposure time was kept low (0.5 or 1 s) and the specimen chamber was moved by 100 μm after each exposure. Following X-ray recordings, background scattering was subtracted, and the major myosin meridional reflection intensities/spacing were determined as described elsewhere previously [84, 85].

### Myosin heavy chain band Isolation for post-translational identifications

Briefly, muscle biopsy samples were cut into 15 mg sections. These were immersed into a sample buffer (0.5 M Tris pH = 6.8, 0.5mg/ml Bromophenol Blue, 10% SDS, 10% Glycerol, 1.25% Mercaptoethanol) at 4°C. Samples were then homogenized and centrifuged allowing the supernatant to be extracted and used for SDS-PAGE gels (with stacking gel made with Acrylamide/Bis 37.5:1 and separation gel made with Acrylamide/Bis 100:1). Proteins were separated and individual MyHC bands excised [86].

### Post-translational Modification Peptide mapping

Proteins were separated on an SDS-PAGE gel (6% polyacrylamide and 30% glycerol), and individual bands excised. For both *U.* arctos and *I.* tridecemlineatus, separate gel bands were excised for Myh7 and Myh2 and their relevant molecular weights. Gel bands were destained twice with destaining buffer (25 mM ammonium bicarbonate, 50% acetonitrile), dehydrated with 100 % acetonitrile, and incubated with reduction/alkylation solution (50 mM Tris pH = 8.5, 10 mM Tcep, 40 mM CAA) for 15 min at 37 °C. Gel bands were washed in destaining buffer, dehydrated, and incubated with 500 ng Trypsin in 20 µL digestion buffer (50 mM TEAB) for 15 min at 37 °C. 30 µL additional digestion buffer was added, and the gel bands incubated over night at 37 °C. After collection of the digested peptides, gel bands were eluted once in 50 µl 1% TFA. Both eluates were combined, and peptides desalted on C18 material prior to LC-MS analysis.

Liquid chromatography was performed using a Vanquish Neo HPLC system (Thermo Fisher Scientific) coupled through a nano-electrospray source to a Tribrid Ascend mass spectrometer (Thermo Fisher Scientific). Peptides were loaded in buffer A (0.1 % formic acid) and separated on a 25 cm column Aurora Gen2, 1.7uM C18 stationary phase (IonOpticks) with a non-linear gradient of 1 – 48% buffer B (0.1 % formic acid, 99.9% acetonitrile) at a flow rate of 400 nL/min over 53 min. The column temperature was kept at 50° C. Spray voltage was set to 2200 V. Data acquisition switched between a full scan (60K resolution, 123ms max. injection time, AGC target 100 %) and 10 data-dependent MS/MS scans (30K resolution, 59 ms max. injection time, AGC target 400 % and HCD activation type). Isolation window was set to 1.4, and normalized collision energy to 25. Multiple sequencing of peptides was minimized by excluding the selected peptide candidates for 45 s.

### Global proteome profiling

Fresh collagenase dilution in DMEM was first prepared by filtering the collagenase through 22 µm. The solution is placed in a +37°C chamber before use. 2 g of each snap frozen sample is added to Eppendorf tubes with marked sample ID. In each tube, 200 μl of +37°C Collagenase is added to break down the connected tissue in the muscle sample to prevent connective tissue in the sample to be analyzed. All tubes were incubated at +37°C for 90 minutes and agitated every 15 min. After this treatment, samples were each put in a 6-well plate. Fibers were then cleaned from connective tissue and separated with tweezers. About 50 fibers of each sample were transferred into an Eppendorf tube with ice-cold PBS. Fibers were spun down at 400g and 4°C and PBS was removed. Skeletal muscle fiber tissue samples were then lysed with lysis buffer (1% (w/v) Sodium Deoxycholate, 100 mM Teab, pH 8.5) and incubated for 10 min at 95°C followed by sonication using a Bioruptor pico (30 cycles, 30 sec on/off, ultra-low frequency). Heat incubation and sonication were repeated once, samples cleared by centrifugation, reduced with 5 mM (final concentration) of TCEP for 15 min at 55°C, alkylated with 20 mM (final concentration) CAA for 30 minutes at RT, and digested adding Trypsin/LysC at 1:100 enzyme/protein ratio. Peptides were cleaned up using StageTips packed with SDB-RPS and resuspended in 50 µL TEAB 100 mM, pH 8,5. 50 µg of each sample was labeled with 0.5 mg TMTpro labeling reagent according to the manufacturer’s instructions. Labeled peptides were combined and cleaned up using C18-E (55 µm, 70 Å, 100 mg) cartridges (Phenomenex).

Labeled desalted peptides were resuspended in buffer A* (5% acetonitrile, 1% TFA), and fractionated into 16 fractions by high-pH fractionation. For this, 20 ug peptides were loaded onto a Kinetex 2.6u EVO C18 100 Å 150 x 0.3 mm column via an EASY-nLC 1200 HPLC (Thermo Fisher Scientific) in buffer AF (10 mM TEAB), and separated with a non-linear gradient of 5 – 44 % buffer BF (10mM TEAB, 80 % acetonitrile) at a flow rate of 1.5 µL / min over 62 min. Fractions were collected every 60 s with a concatenation scheme to reach 16 final fractions (e.g. fraction 17 was collected together with fraction 1, fraction 18 together with fraction 2, and so on).

Fractions were evaporated, resuspended in buffer A*, and measured on a Vanquish Neo HPLC system (Thermo Fisher Scientific) coupled through a nano-electrospray source to a Tribrid Ascend mass spectrometer (Thermo Fisher Scientific). Peptides were loaded in buffer A (0.1 % formic acid) onto a 110 cm mPAC HPLC column (Thermo Fisher Scientific) and separated with a non-linear gradient of 1 – 50 % buffer B (0.1 % formic acid, 80 % acetonitrile) at a flow rate of 300 nL/min over 100 min. The column temperature was kept at 50° C. Samples were acquired using a Real Time Search (RTS) MS3 data acquisition where the Tribrid mass spectrometer was switching between a full scan (120 K resolution, 50 ms max. injection time, AGC target 100%) in the Orbitrap analyzer, to a data-dependent MS/MS scans in the Ion Trap analyzer (Turbo scan rate, 23 ms max. injection time, AGC target 100% and HCD activation type). Isolation window was set to 0.5 (m/z), and normalized collision energy to 32. Precursors were filtered by charge state of 2-5 and multiple sequencing of peptides was minimized by excluding the selected peptide candidates for 60 s. MS/MS spectra were searched in real time on the instrument control computer using the Comet search engine with either the UP000291022 U. americanus or UP000005215 *I. tridecemlineatus* FASTA file, 0 max miss cleavage, 1 max oxidation on methionine as variable mod. and 35 ms max search time with an Xcorr soring threshold of 1.4 and 20 precursor ppm error). MS/MS spectra resulting in a positive RTS identification were further analyzed in MS3 mode using the Orbitrap analyzer (45K resolution, 105 ms max. injection time, AGC target 500%, HCD collision energy 55 and SPS = 10). The total fixed cycle time, switching between all 3 MS scan types, was set to 3 s.

### Proteomics Data analysis

Raw mass spectrometry data from peptide mapping experiments were analyzed with MaxQuant (v2.1.4). Peak lists were searched against the *U. americanus* (Uniprot UP000291022) or Ictidomys tridecemlineatus (Uniprot UP000005215) proteomes combined with 262 common contaminants by the integrated Andromeda search engine. False discovery rate was set to 1 % for both peptides (minimum length of 7 amino acids) and proteins. Phospho(STY) and Acetyl(K) were selected as variable modifications.

Raw mass spectrometry data from global proteome profiling were analyzed with Proteome Discoverer (v3.0.1.27) using the default processing workflow “PWF_Tribrid_TMTpro_SPS_MS3_SequestHT_INFERYS_Rescoring_Percolator”. Briefly, peak lists were searched against the UniProtKB UP000291022 *U. americanus* or UP000005215 *I. tridecemlineatus* FASTA databases by the integrated SequestHT search engine, setting Carbamidomethyl (C) and TMTpro (K, N-Term) as static modifications, Oxidation (M) as variable modification, max missed cleavage as 2 and minimum peptide amino acid length as 7. The false discovery rate was set to 0.01 (strict) and 0.05 (relaxed).

All statistical analysis of TMT derived protein expression data was performed using in-house developed python scripts based on the analysis pipeline of the Clinical Knowledge Graph [87]. Protein abundances were log2-transformed, and proteins with less than 2 valid values in at least one group were excluded from the analysis. Missing values were imputed with MinProb approach (width=0.3 and shift=1.8). Statistically significant proteins were determined by unpaired t-tests, with Benjamini-Hochberg correction for multiple hypothesis testing. Fold-change (FC) and False Discovery Rate (FDR) thresholds were set to 2 and 0.05 (5%), respectively.

The mass spectrometry proteomics data have been deposited to the ProteomeXchange Consortium via the PRIDE partner repository [88] with the dataset identifier PXD044505, PXD044685 and PXD044728.

Gene Ontology networks were created using Metascape software [89]. Networks were annotated using Cytoscape [90]. Principal component plots analysis were created in Perseus [91].

### 1H-NMR Metabolomic characterization

Frozen lyophilized tissue samples were shipped on dry ice to Biosfer Teslab (Reus, Spain) for the ^1^H-NMR analysis. Prior to analysis, aqueous and lipid extracts were obtained using the Folch method with slight modifications [92]. Briefly, 1440 µL of dichloromethane:methanol (2:1, v/v) were added to 25 mg of pulverized tissue followed by three 5 min sonication steps with one shaking step in between. Next, 400 µL of ultrapure water was added, mixed, and centrifuged at 25100 g during 5 min at 4 °C. Aqueous and lipid extracts were transferred to a new Eppendorf tube and completely dried in SpeedVac to achieve solvent evaporation and frozen at -80 °C until ^1^H-NMR analysis.

Aqueous extracts were reconstituted in a solution of 45 mM PBS containing 2.32 mM of Trimethylsilylpropanoic acid (TSP) as a chemical shift reference and transferred into 5-mm NMR glass tubes. ^1^H-NMR spectra were recorded at 300 K operating at a proton frequency of 600.20 MHz using an Avance III-600 Bruker spectrometer. One-dimensional ^1^H pulse experiments were carried out using the nuclear Overhauser effect spectroscopy (NOESY)-presaturation sequence to suppress the residual water peak at around 4.7 ppm and a total of 64k data points were collected. The acquired spectra were phased, baseline-corrected and referenced before performing the automatic metabolite profiling of the spectra datased through and adaptation of Dolphin [93]. Several database engines (Bioref AMIX database (Bruker), Chenomx and HMDB, and literature were used for 1D-resonances assignment and metabolite identification [94, 95].

Lipid extracts were reconstituted in a solution of CDCl_3_:CD_3_OD:D_2_O (16:7:1, v/v/v) containing Tetramethylsilane (TMS) and transferred into 5-mm NMR glass tubes. ^1^H-NMR spectra were recorded at 286 K operating at a proton frequency of 600.20 MHz using an Avance III-600 Bruker spectrometer. A 90° pulse with water pre-saturation sequence (ZGPR) was used. Quantification of lipid signals in 1H-NMR spectra was carried out with LipSpin an in-house software based on Matlab [96]. Resonance assignments were done based on literature values [95].

### Myosin Chimera Simulation

ChimeraX was used to make a simulation of Myh2 to illustrate the changes occurring during torpor in *I. tridecemlineatus*. The sequence was downloaded from UniProt, and significant post-translational modifications positions were highlighted on the protein simulation and marked with red. ATP binding domain and actin binding domain were additionally highlighted.

### EvoEF Protein Stability Simulations

EvoEF (version 1) was used to calculate the stability change upon mutation, in terms of ΔΔG. To this end, we first used "EvoEF --command=RepairStructure" to repair clashes and torsional angles of the wild type structure. "EvoEF --command=BuildMutant" is then used to mutate the repaired wild type structures into the mutant by changing the side chain amino acid type followed by a local side chain repacking. "EvoEF --command=ComputeStability" is then applied to both the repair wild type and the mutant to calculate their respective stabilities (ΔGWT and ΔGmutant). The stability change upon mutation can then be derived by ΔΔG=ΔGmutant-ΔGWT. A ΔΔG below zero means that the mutation causes destabilization; otherwise, it induces stabilization [35, 97]. The sequence of the MYH2 coiled-coil backbone was used for EvoEF stability calculations due to size limitations in the software when using the entire MYH2 protein sequence.

### Statistical analysis

Data are presented as means ± standard deviations. Statistical tests used are listed in the figure legends.Graphs were prepared and analysed in Graphpad Prism v9. Statistical significance was set to *p* < 0.05 unless otherwise stated.

## Acknowledgements

This work was generously funded by the Carlsberg Foundation (CF20-0113) grant to J.O. The X-ray experiments were performed under approval of the SPring-8 Proposal Review Committee (2022A1069 and 2022B1107). The related X-ray data reduction and analyses were performed by Accelerated Muscle Biotechnologies Consultants LLC (USA). Mass spectrometry analyses were performed by the Proteomics Research Infrastructure (PRI) at the University of Copenhagen, supported by the Novo Nordisk Foundation (grant agreement number NNF19SA0059305). The Scandinavian brown bear research project is funded by the Norwegian Environment Agency and the Swedish Environmental Protection Agency. KLD, AVG and VBF were supported by P20GM130443. Tissue collection from I. tridecemlineatus was supported by a Discovery Grant to JFS from the Natural Sciences and Engineering Research Council (Canada).

## Author Contributions

JO acquired funding; CTAL and JO conceived the study; CTAL and JO managed the project; CTAL, EGM, MMO, MSO, JL, RAES, MNG, CZ, HI, ALH, MNK, CM, NA, OF, SG, JFS, AVG, VBF, BMB, ØT and KLD performed experiments; CTAL, EGM, MMO, MSO, JL, RAES, MNG, CZ, HI, ALH, MNK, CM, NA, OF, SG, RJS, JFS and JO analysed data and interpreted the results; CTAL, EGM, MMO, MSO, JL, RAES, MNG, CZ, HI, ALH, MNK, CM, NA, OF, SG, JFS, AVG, VBF, BMB, ØT, KLD and JO wrote and reviewed the manuscript.

