## Supplementary Figures and Tables for "Remodelling of skeletal muscle myosin metabolic states in hibernating mammals"

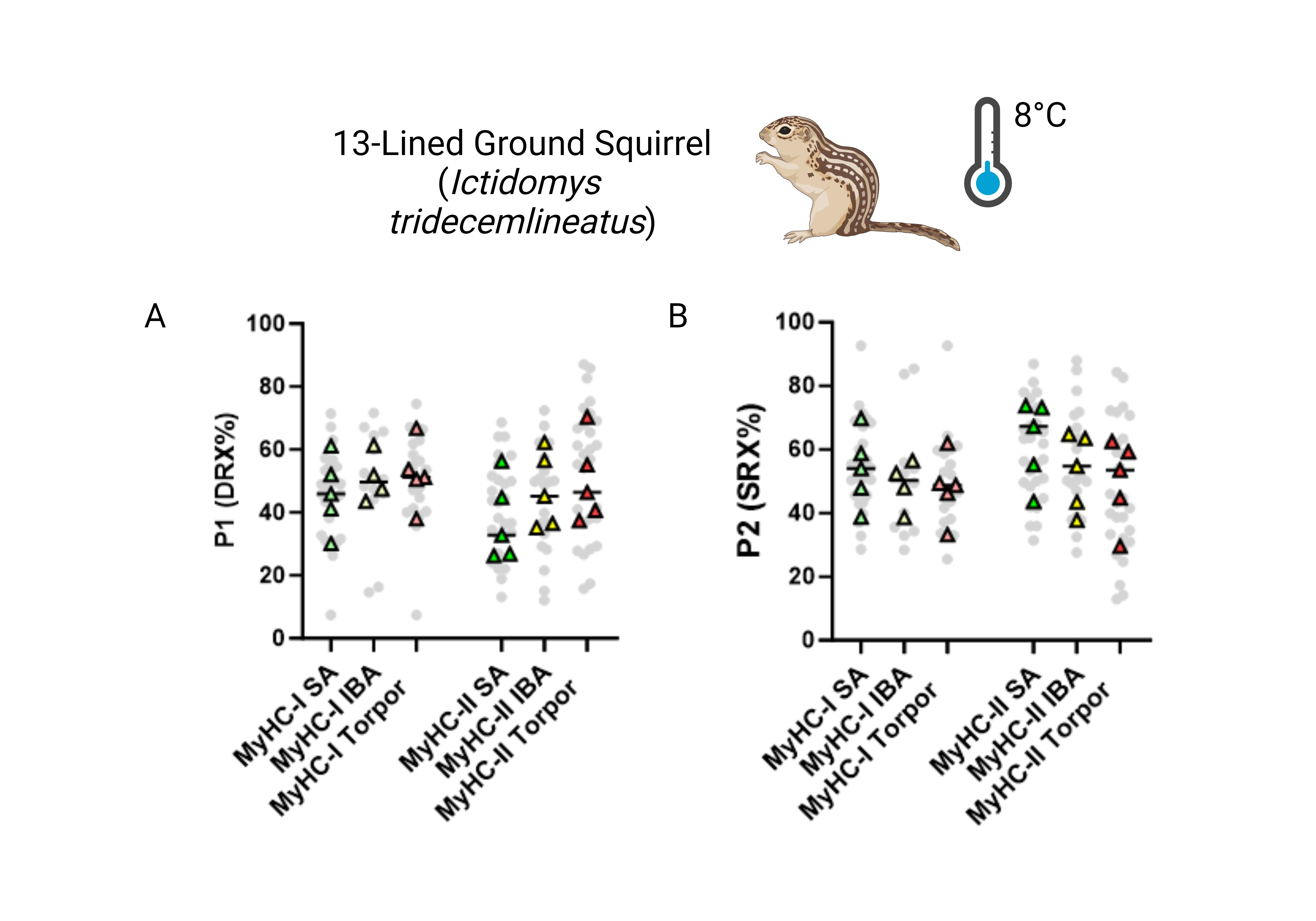

**Supplementary Figure 1 – Myosin ATP turnover lifetime is altered following exposure to cold temperature in MyHC-II muscle fibers from *I. tridecemlineatus* during active and IBA periods but not torpor. A-B.** Percentage of myosin heads in the P1/DRX (A) or P2/SRX (B) from *I. tridecemlineatus* single muscle fibers obtained during summer active (SA), interbout arousal (IBA) or torpor periods at 8°C. Grey circles represent the values from each individual muscle fiber which was analyzed. Colored triangles represent the mean value from an individual animal, 8-12 fibers analyzed per animal. Statistical analysis was performed upon mean values. One-way ANOVA was used to calculate statistical significance. n = 5 individual animals per group.

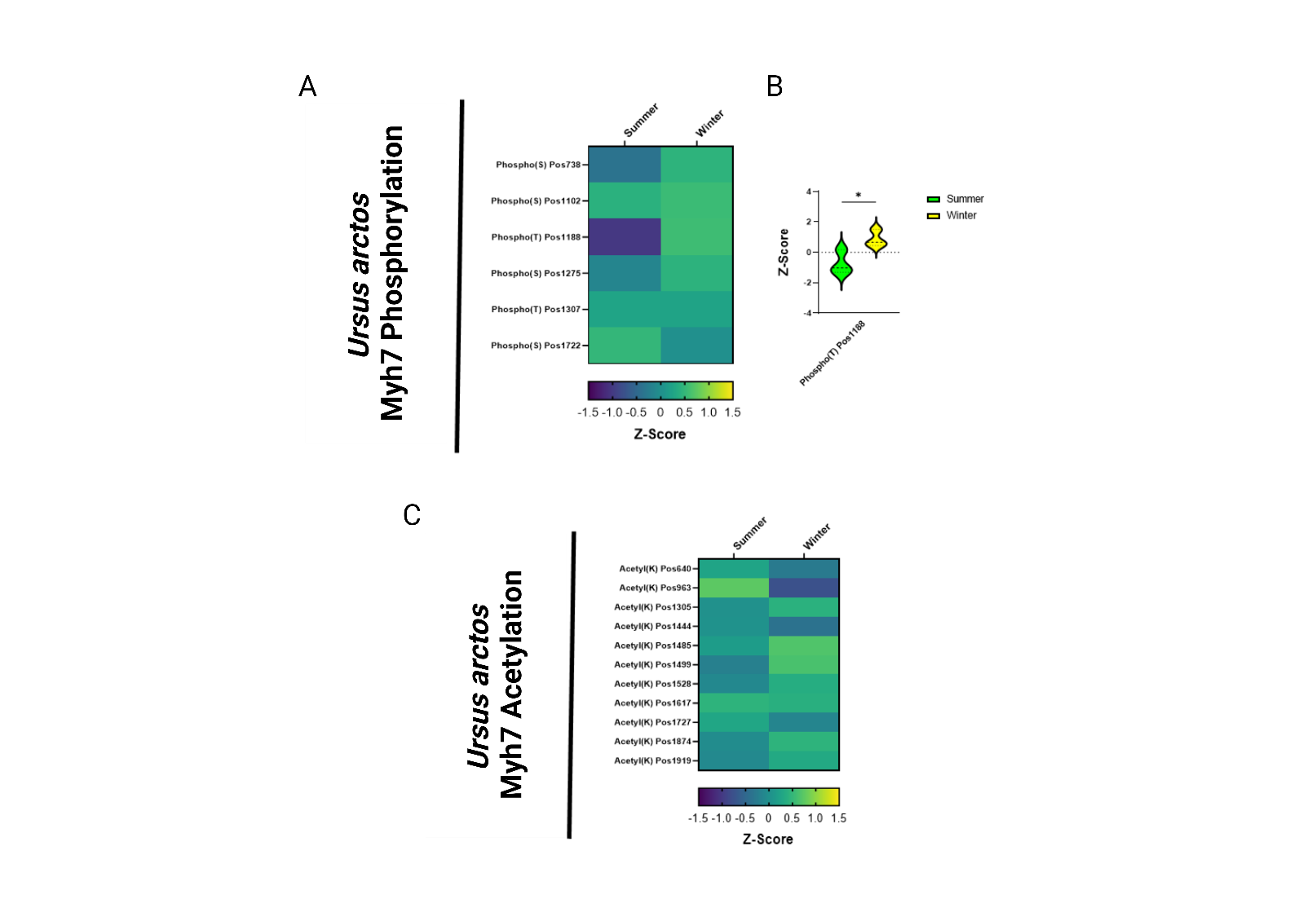

**Supplementary Figure 2 – MYH7 protein phosphorylation and acetylation in *U. arctos* is relatively unchanged during winter periods. A.** Peptide mapping of differentiated phosphorylation sites upon MYH7 protein during summer and winter periods. Heat map demonstrates all sites observed to be differentiated following the calculation of z-scores for each site. Z-scores > 0 equal hyper-phosphorylation and z-scores < 0 equal hypo-phosphorylation for each residue. **B.** Violin plot demonstrates significantly differentiated residue using z-scores. **C.** Peptide mapping of differentiated acetylation sites upon MYH7 protein during summer and winter periods. Heat map demonstrates all sites observed to be differentiated following the calculation of z-scores for each site. Z-scores > 0 equal hyper-acetylation and z-scores < 0 equal hypo-acetylation for each residue. Two-way ANOVA with Šídák's multiple comparisons test was used to calculate statistical significance. * = *p* < 0.05. n = 5 individual animals per group.

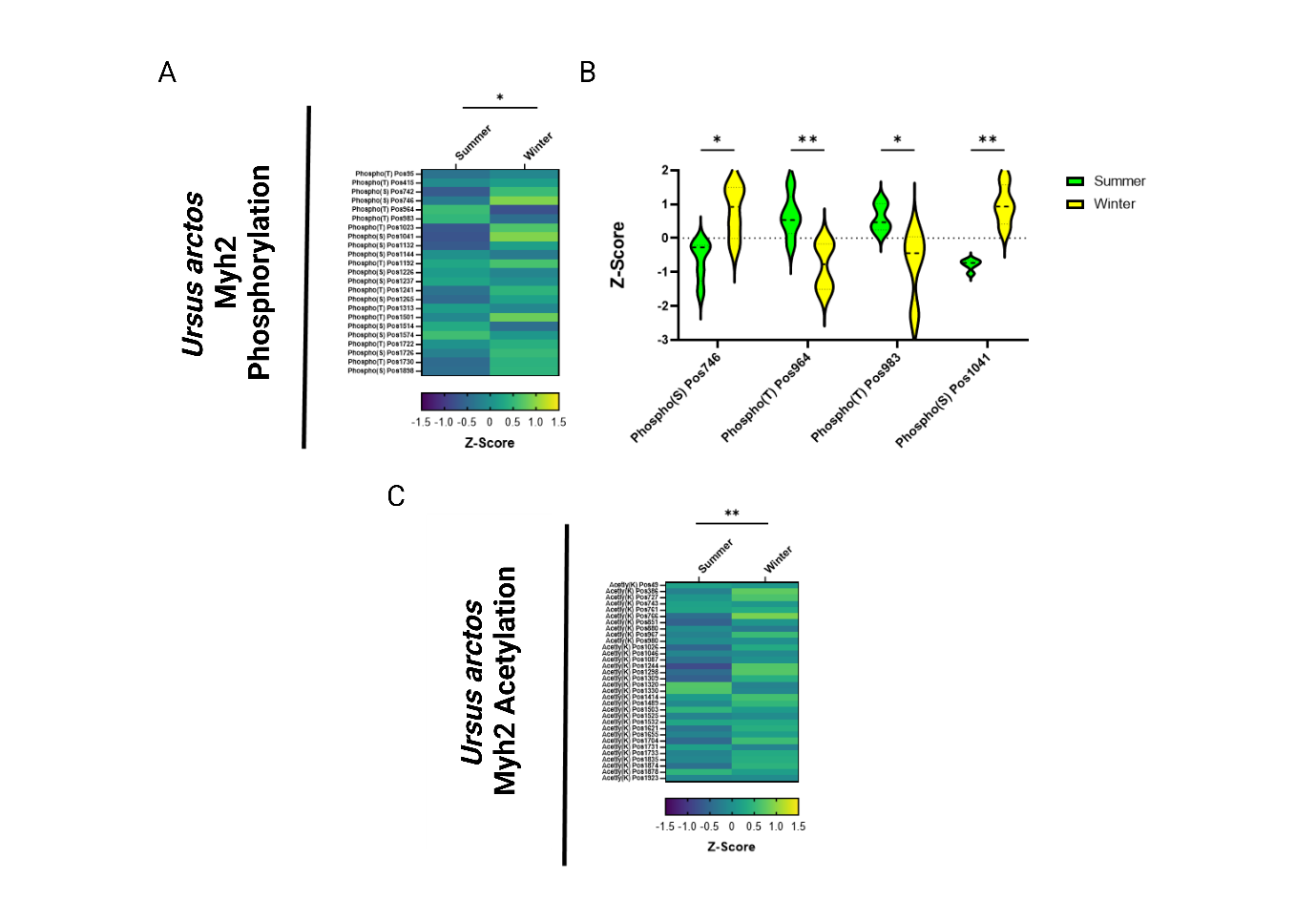

**Supplementary Figure 3 – MYH2 protein phosphorylation and acetylation in *U. arctos* is altered during winter periods. A.** Peptide mapping of differentiated phosphorylation sites upon MYH2 protein during summer and winter periods. Heat map demonstrates all sites observed to be differentiated following the calculation of z-scores for each site. Z-scores > 0 equal hyper-phosphorylation and z-scores < 0 equal hypo-phosphorylation for each residue. **B.** Violin plot demonstrates significantly differentiated residues using z-scores. **C.** Peptide mapping of differentiated acetylation sites upon MYH7 protein during summer and winter periods. Heat map demonstrates all sites observed to be differentiated following the calculation of z-scores for each site. Z-scores > 0 equal hyper-acetylation and z-scores < 0 equal hypo-acetylation for each residue. Two-way ANOVA with Šídák's multiple comparisons test was used to calculate statistical significance. * = *p* < 0.05, ** = *p* < 0.01. n = 5 individual animals per group.

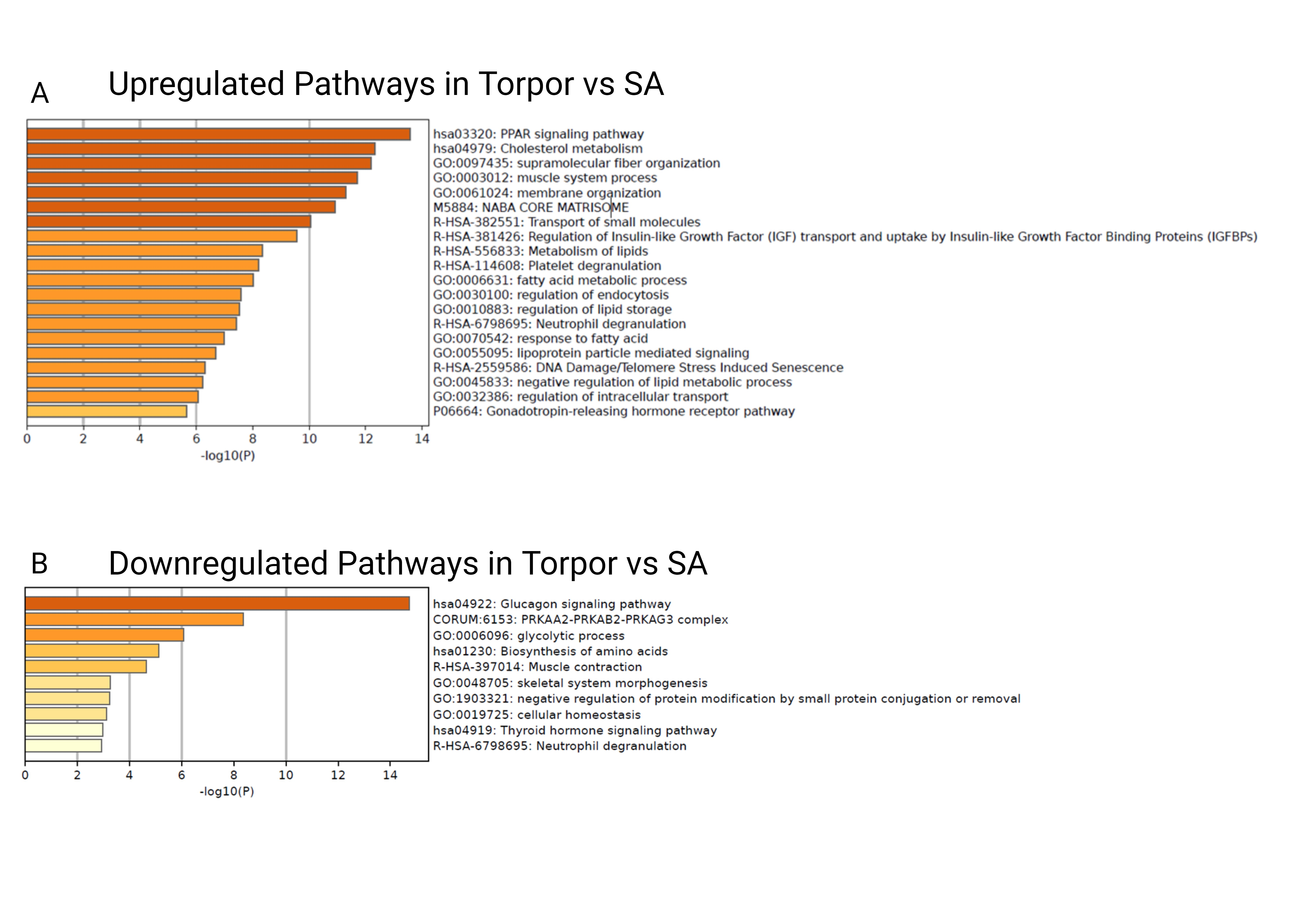

**Supplementary Figure 4 – All ontological clusters altered in *I. tridecemlineatus* in torpor vs SA periods. A.** All upregulated pathways identified in Metascape in torpor vs SA periods. Pathways are listed in order of significance by -log10(*p*). **B.** All downregulated pathways identified in Metascape in torpor vs SA periods. Pathways are listed in order of significance by -log10(*p*).

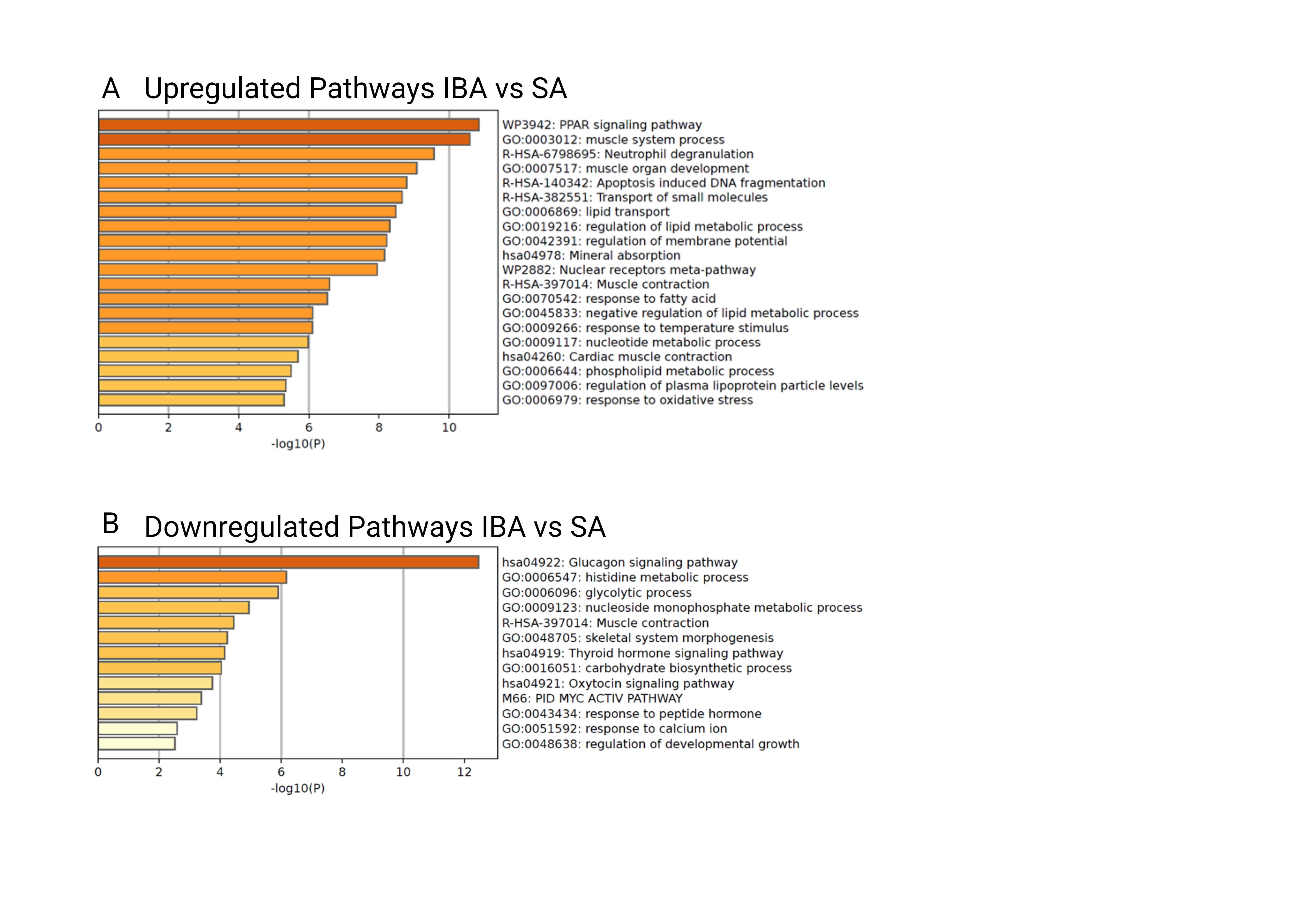

**Supplementary Figure 5 – All ontological clusters altered in *I. tridecemlineatus* in IBA vs SA periods. A.** All upregulated pathways identified in Metascape in IBA vs SA periods. Pathways are listed in order of significance by -log10(*p*). **B.** All downregulated pathways identified in Metascape in IBA vs SA periods. Pathways are listed in order of significance by -log10(*p*).

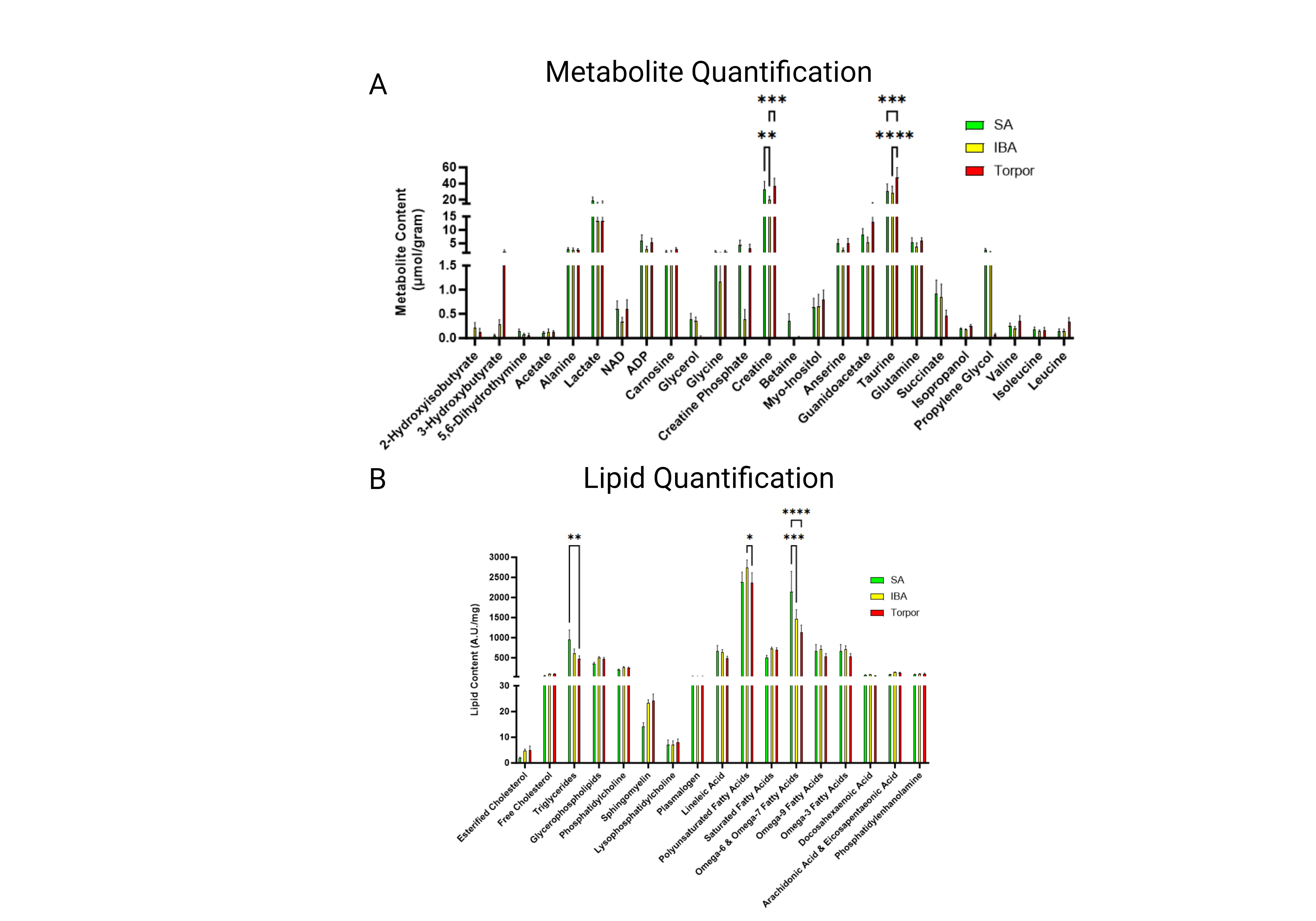

**Supplementary Figure 6 – Metabolite and Lipid Quantification of skeletal muscle from *I. tridecemlineatus* reveals a decrease in lipid levels during torpor. A.** Quantification of metabolites from the skeletal muscle tissue of *I. tridecemlineatus* during SA, IBA and torpor periods. Presented in µmol of metabolite per gram of tissue. **B.** Quantification of lipids from the skeletal muscle tissue of *I. tridecemlineatus* during SA, IBA and torpor periods. Presented in A.U. of lipid per mg of tissue. Two-way ANOVA with Turkey’s multiple comparisons test was used to calculate statistical significance. * = *p* < 0.05, ** = *p* < 0.01, *** = *p* < 0.001, **** = *p* < 0.0001. n=5 individual animals per group.

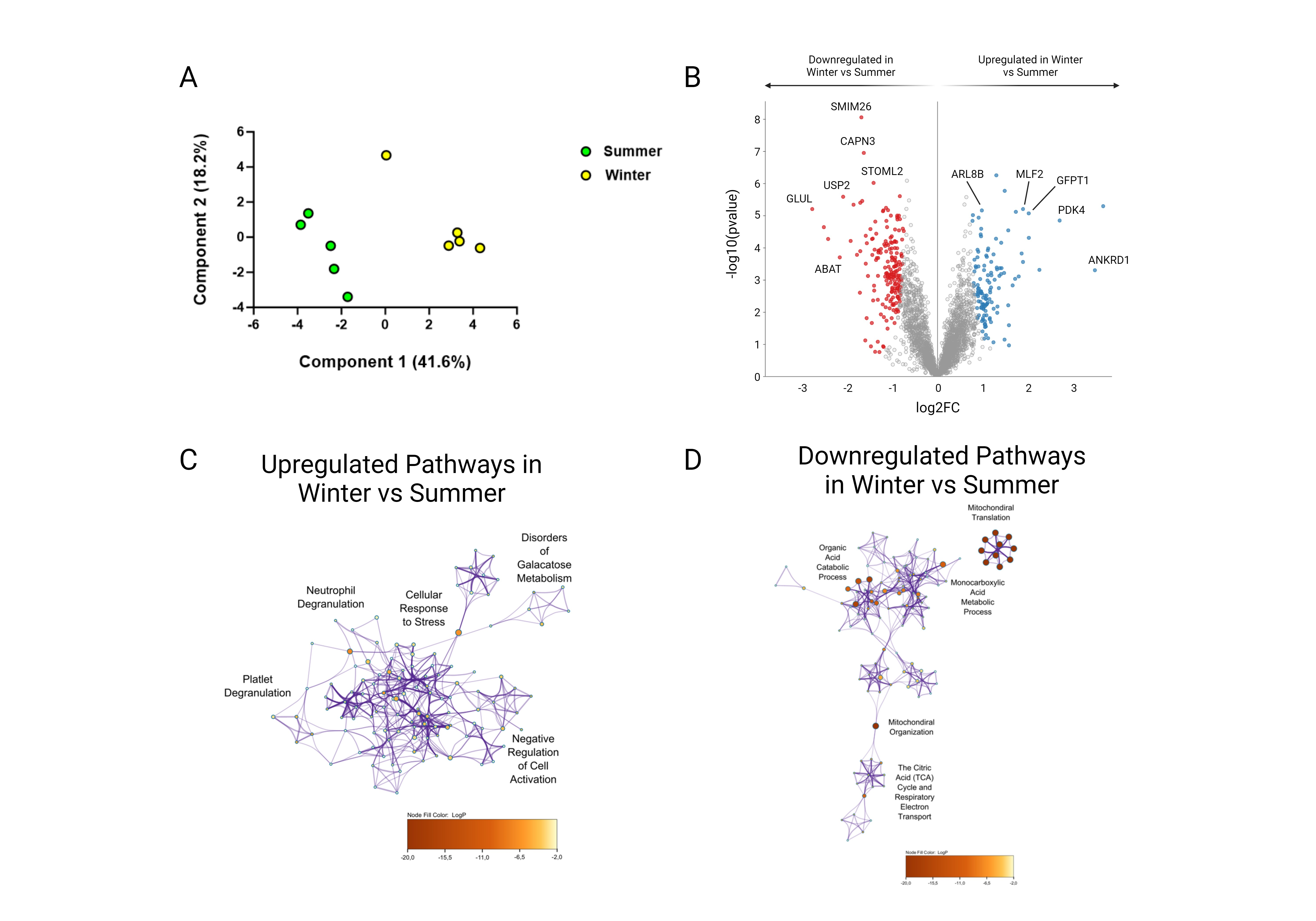

**Supplementary Figure 7 – Global Proteome Analysis of *U. arctos* skeletal muscle fibers reveal metabolic changes but not sarcomeric changes. A.** Principal component analysis for all animals analyzed during summer and winter periods. **B.** Volcano plot displaying proteins which are differentially expressed during winter vs summer periods. FDR < 0.01. Red circles are upregulated proteins and blue circles are downregulated proteins. Highly differentiated proteins of interest are annotated with their respective protein name. **C.** Ontological associations between proteins upregulated during winter vs summer periods. The top five association clusters are annotated on the network. A full list of clusters and the proteins lists included in clusters are available in supplementary figure 4 and supplementary table 4. **D.** Ontological associations between proteins downregulated during winter vs summer periods. The top five association clusters are annotated on the network. A full list of clusters and the proteins lists included in clusters are available in supplementary figure 4 and supplementary table 4. Gene ontology networks were established using Metascape and visualized using Cytoscape. Detailed information of the statistical testing used is available in the methods section. FDR < 0.01 significantly differentially expressed proteins were used to establish networks. Purple lines indicate a direct interaction. Circle size is determined by enrichment and color is determined by *p* value. n = 5 individual animals per group.

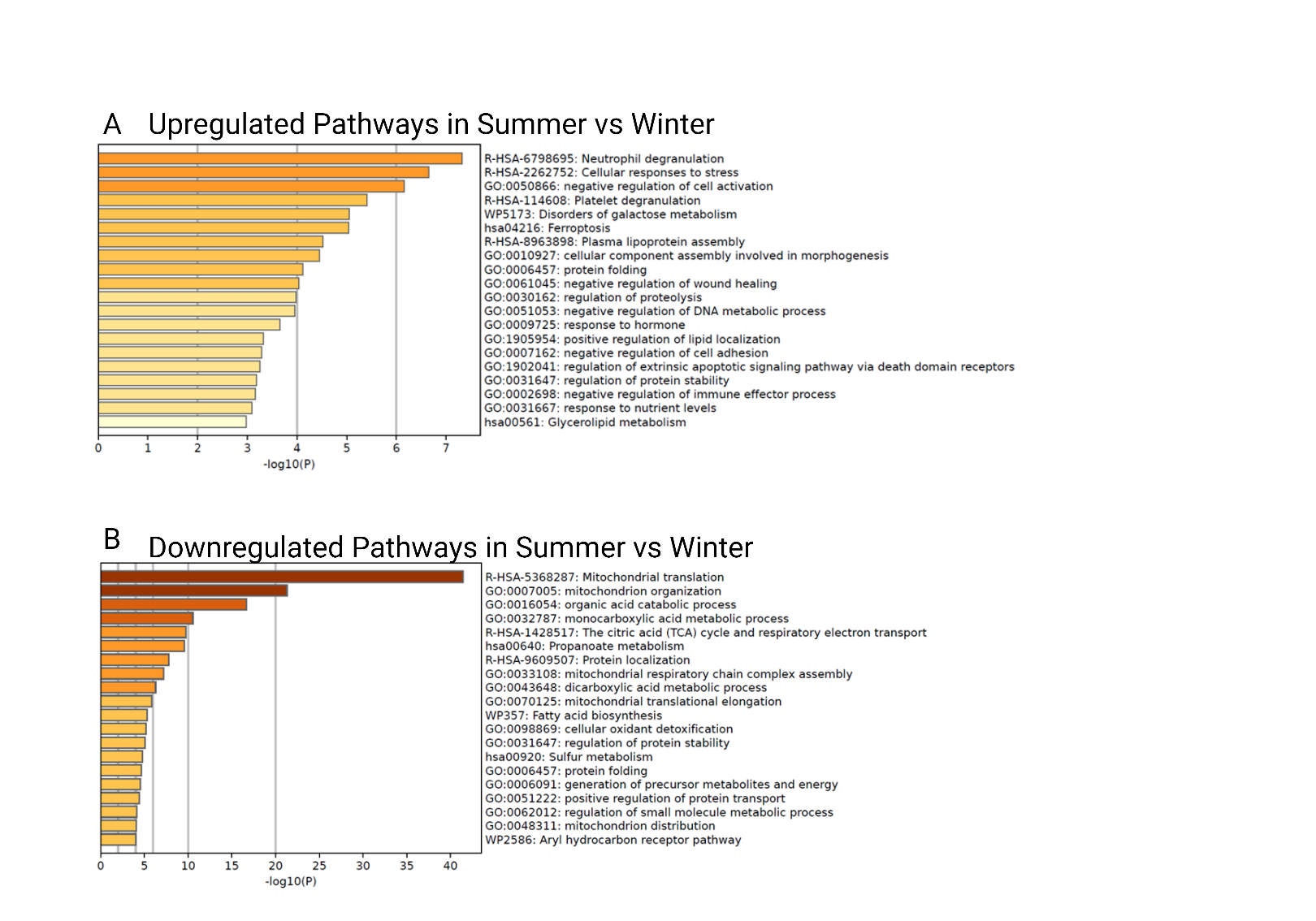

**Supplementary Figure 8 – All ontological clusters altered in *U. arctos* in summer vs winter periods. A.** All upregulated pathways identified in Metascape in winter vs summer periods. Pathways are listed in order of significance by -log10(*p*). **B.** All downregulated pathways identified in Metascape in winter vs summer periods. Pathways are listed in order of significance by -log10(*p*).

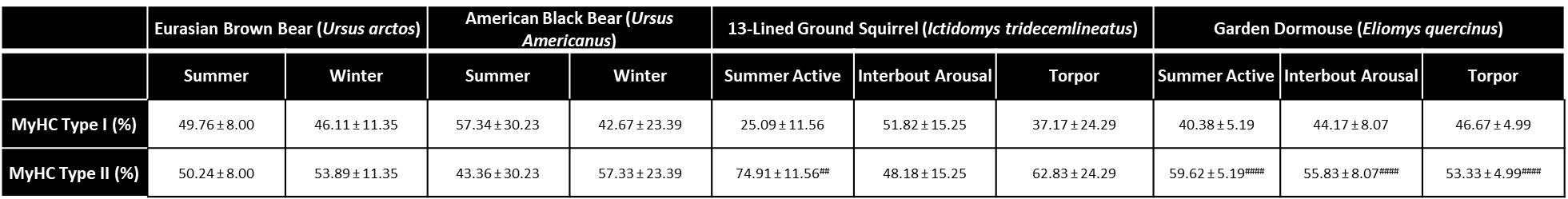

**Supplementary Table 1 – Fiber type compositions from animals used for this study.** Table demonstrating the percentage of fibers which were analyzed during Mant-ATP chase assays that were either MyHC type I or MyHC type II. Data is presented as mean for each animal ± SD. One-way ANOVA was used to calculate significance between hibernation periods in *I. tridecemlineatus* and *E. quercinus*. Student’s t-test was used to calculate significant between hibernating periods in *U.* arctos and *U. americanus* and between MyHC type I and MyHC type II in all animals. ## = *p* < 0.01 vs MyHC type I in corresponding group. #### = *p* < 0.0001 vs MyHC type I in corresponding group. n = 5 individual animals per group.

A

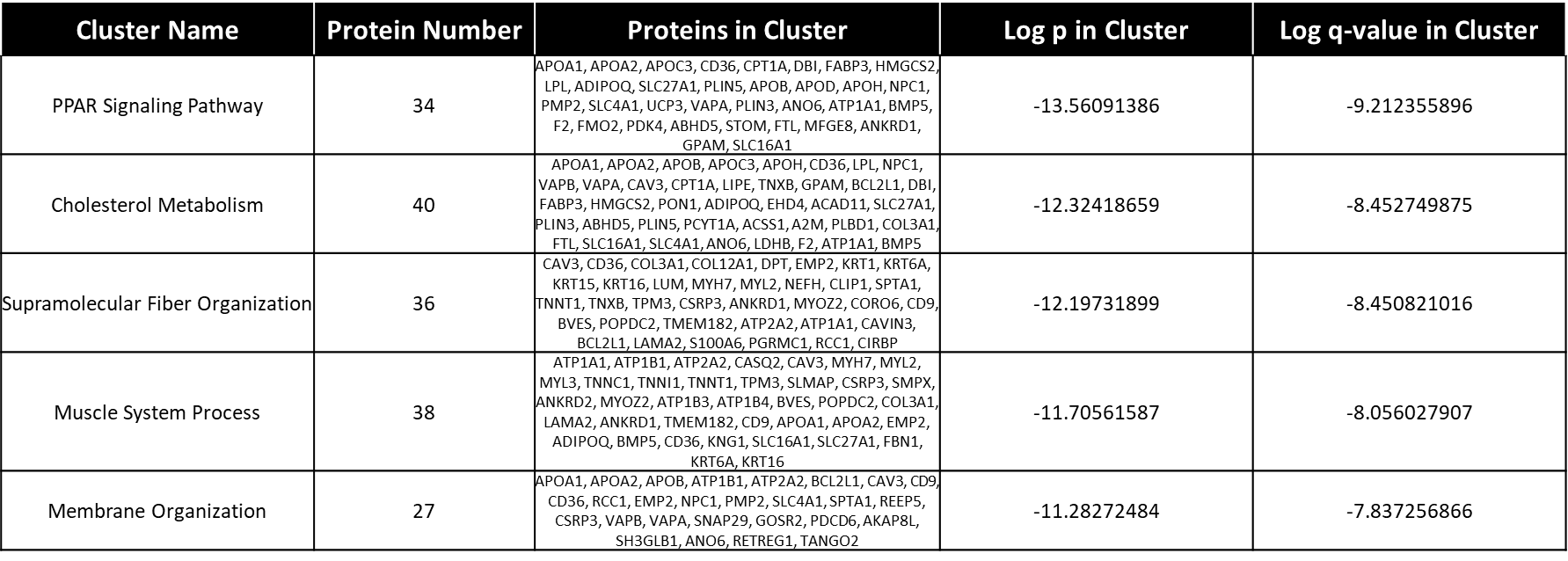

B

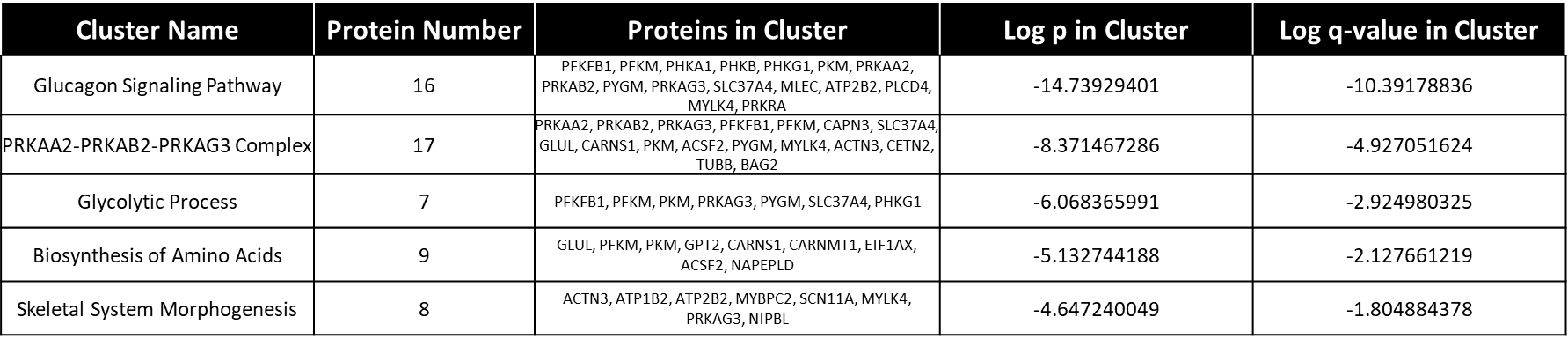

**Supplementary Table 2 – Detailed protein lists for top five differentially expressed ontological clusters in *I. tridecemlineatus* in torpor vs SA periods. A.** Table details the list of proteins which were found to be differentially upregulated in torpor vs SA in each corresponding ontological cluster as identified by Metascape. Clusters are arranged by order of statistical significance. Proteins are listed in alphabetical order within each cluster. **B.** Table details the list of proteins which were found to be differentially downregulated in torpor vs SA in each corresponding ontological cluster as identified by Metascape. Clusters are arranged by order of statistical significance. Proteins are listed in alphabetical order within each cluster. n = 5 individual animals per group.

A

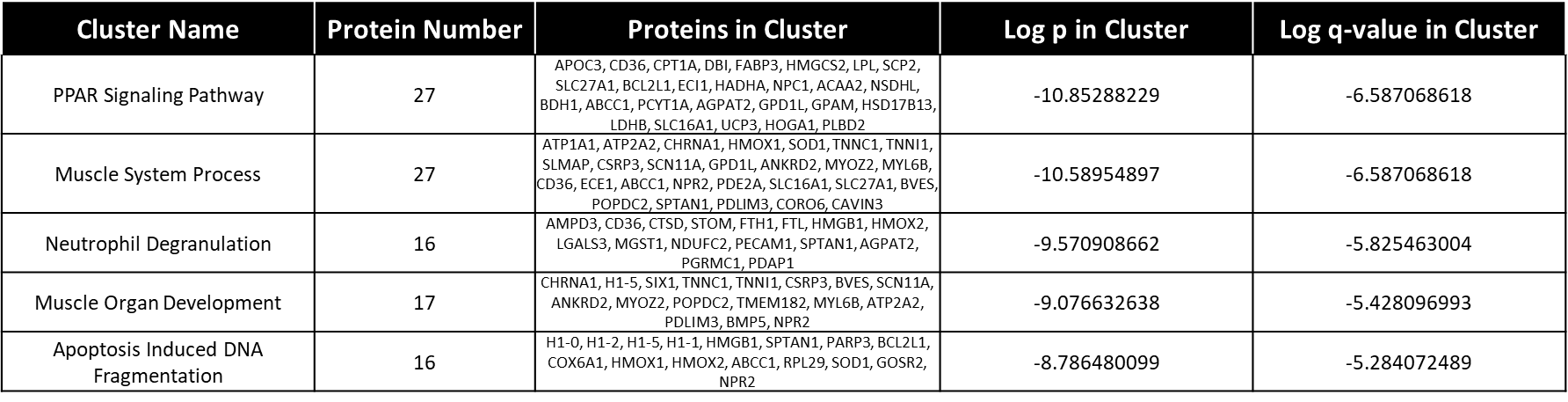

B

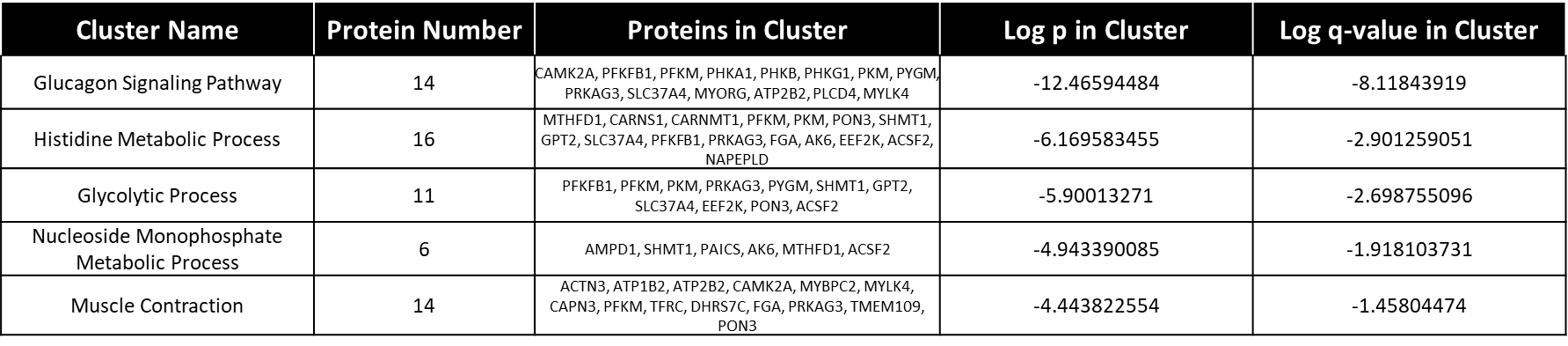

**Supplementary Table 3 – Detailed protein lists for top five differentially expressed ontological clusters in *I. tridecemlineatus* in IBA vs SA periods. A.** Table details the list of proteins which were found to be differentially upregulated in IBA vs SA in each corresponding ontological cluster as identified by Metascape. Clusters are arranged by order of statistical significance. Proteins are listed in alphabetical order within each cluster. **B.** Table details the list of proteins which were found to be differentially downregulated in IBA vs SA in each corresponding ontological cluster as identified by Metascape. Clusters are arranged by order of statistical significance. Proteins are listed in alphabetical order within each cluster. n = 5 individual animals per group.

A

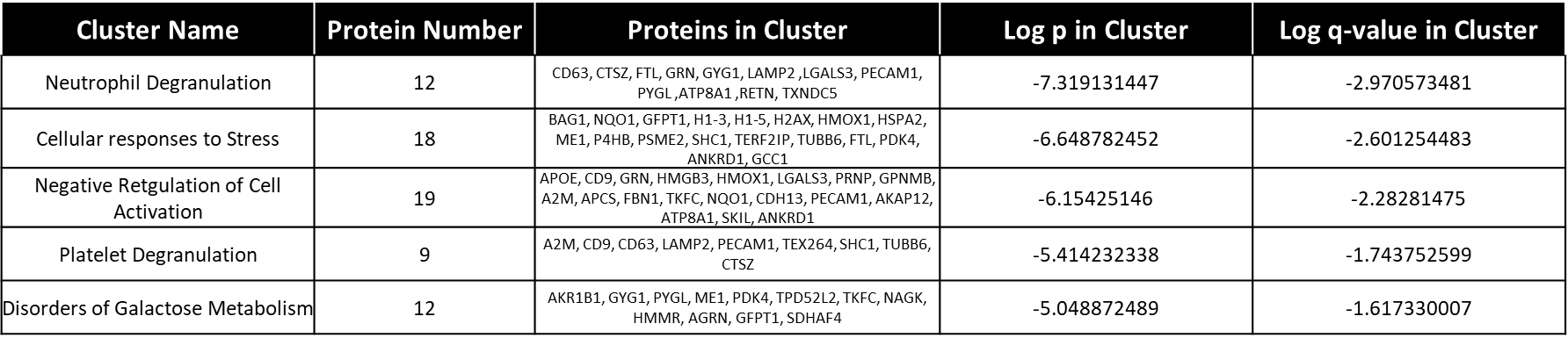

B

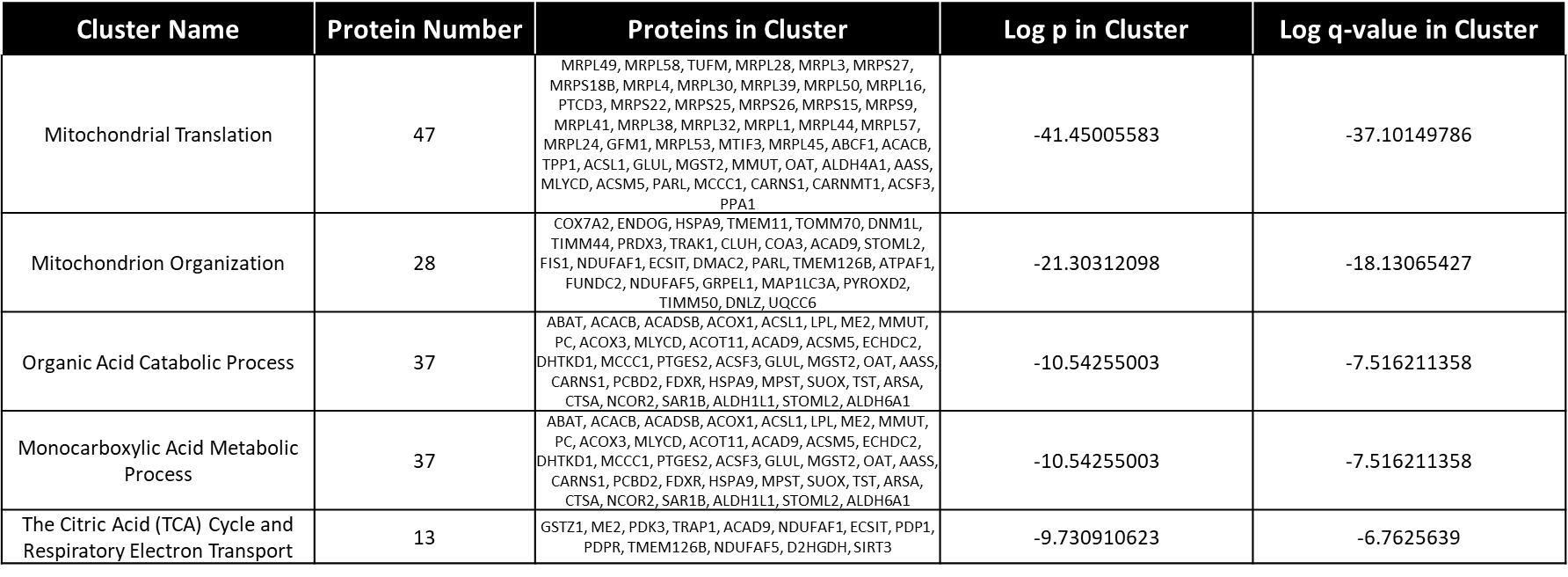

**Supplementary Table 4 – Detailed protein lists for top five differentially expressed ontological clusters in *U. arctos* in winter vs summer periods. A.** Table details the list of proteins which were found to be differentially upregulated in winter vs summer in each corresponding ontological cluster as identified by Metascape. Clusters are arranged by order of statistical significance. Proteins are listed in alphabetical order within each cluster. **B.** Table details the list of proteins which were found to be differentially downregulated in winter vs summer in each corresponding ontological cluster as identified by Metascape. Clusters are arranged by order of statistical significance. Proteins are listed in alphabetical order within each cluster. n = 5 individual animals per group.
